# MicroRNA-143 plays a protective role in ischemia-induced retinal neovascularization

**DOI:** 10.1101/548297

**Authors:** Jiang-Hui Wang, Jinying Chen, Damien Ling, Leilei Tu, Vikrant Singh, Moeen Riaz, Fan Li, Selwyn M. Prea, Zheng He, Bang V. Bui, Alex W. Hewitt, Peter van Wijngaarden, Gregory J. Dusting, Guei-Sheung Liu

**Affiliations:** Centre for Eye Research Australia, Royal Victorian Eye and Ear Hospital, East Melbourne, Victoria, Australia; Ophthalmology, Department of Surgery, University of Melbourne, East Melbourne, Victoria, Australia; Menzies Institute for Medical Research, University of Tasmania, Hobart, Tasmania, Australia; Department of Ophthalmology, the First Affiliated Hospital of Jinan University, Guangzhou, Guangdong, China; Discipline of Ophthalmology, Sydney Medical School, University of Sydney, Sydney, New South Wales, Australia; Public Health Genomics, School of Public Health and Preventive Medicine, Monash University, Melbourne, Victoria, Australia.; State Key Laboratory of Ophthalmology, Zhongshan Ophthalmic Centre, Sun Yat-sen University, Guangzhou, Guangdong, China; Department of Optometry and Vision Sciences, University of Melbourne, Parkville, Victoria, Australia

**Keywords:** MicroRNA, Next-generation sequencing, Retinal neovascularization

## Abstract

Retinal neovascularization is a severe complication of proliferative diabetic retinopathy. MicroRNAs (miRNAs) are master regulators of gene expression that play important roles in retinal neovascularization. Here, we investigated the retinal miRNA expression profile in a rat model of oxygen-induced retinopathy (OIR) through miRNA-Seq. We found that miR-143-3p, miR-126-3p, miR-150-5p and miR-145-5p were significantly down-regulated in the retina of OIR rats, and directly involved in the development of retinal neovascularization. Of these identified miRNAs, miR-143 is enriched in retina and was first reported being associated with pathological retinal angiogenesis. Our RNA-Seq data further suggested that miR-143 alleviates retinal neovascularization by mediating the inflammation/stress pathways via *Fos*. Moreover, the computational analysis indicated that Transforming Growth Factor-beta Activated Kinase 1 (*TAK1*) is involved in several key pathways associated with the dysregulated miRNAs. The pharmacological inhibition of *TAK1* suppressed angiogenesis *in vitro* and retinal neovascularization *in vivo*. Our data highlight the utility of next-generation sequencing in the development of therapeutics for ocular neovascularization and further suggest that therapeutic targeting the dysregulated miRNAs or TAK1 may be a feasible adjunct therapeutic approach in patients with retinal neovascularization.

## Introduction

Retinal neovascularization is a sight-threatening sequela of several common diseases, including proliferative diabetic retinopathy (1). The development of abnormal, new blood vessels may lead to serious complications that cause vision loss and blindness, such as vitreous hemorrhage and retinal detachment. Although the increased understanding of the pathogenesis of retinal angiogenesis has led to the development of effective treatments, most notably vascular endothelial growth factor (VEGF) inhibitors, certain patients respond poorly to treatment, indicating that other pathways may be involved. Furthermore, the cost-effectiveness of the prolonged use of VEGF inhibitors (e.g., ranibizumab and aflibercept) may have deleterious consequences (2). Thus, novel therapeutic options for patients with retinal neovascularization are needed.

MicroRNAs (miRNAs) constitute a class of endogenous non-coding RNA molecules of approximately 22 nucleotides in length. miRNAs act by complementary base pairing at the 3’ untranslated region (3’ UTR) of target mRNA, resulting in the degradation or translational repression of the target mRNA (3). miRNAs play important roles in several biological processes, including cellular growth, development, differentiation, homeostasis, and death (4). A single miRNA typically has hundreds of potential target mRNAs, and conversely, a single mRNA may be targeted by multiple miRNAs (3). miRNAs have been shown to be important modulators of blood vessel formation and function. The dysregulation of retinal miRNAs, such as miRNA-150 (miR-150), miRNA-126 (miR-126), and miRNA-155 (miR-155), may cause the progression of retinal neovascularization in animal models (5-7). However, these studies have yielded inconsistent results, which are likely due to different protocols of establishing animal models. Moreover, the potential of using these dysregulating miRNAs as a treatment for retinal neovascularization is unclear.

In this study, we performed next-generation sequencing (NGS) to identify the dysregulated miRNAs in a rat model of oxygen-induced retinopathy (OIR) (Fig. 1A). Next, we evaluated the involvement of these dysregulated miRNAs in the development of retinal neovascularization in the OIR rats (Fig. 1B). Finally, we interrogated the putative functions of the dysregulated miRNAs through the *in-silico* analysis and identified transforming Growth Factor-beta Activated Kinase 1 (*TAK1*), which is a serine/threonine kinase, was involved in retinal neovascularization. Selective inhibition of TAK1 by 5Z-7-oxozeaenol was showed to suppress angiogenesis *in vitro* and *in vivo* (Fig. 1C). Our study highlights the utility of computational analyses using NGS data in screening for novel genes involved in complex pathological processes, such as retinal neovascularization.

**Figure 1.**
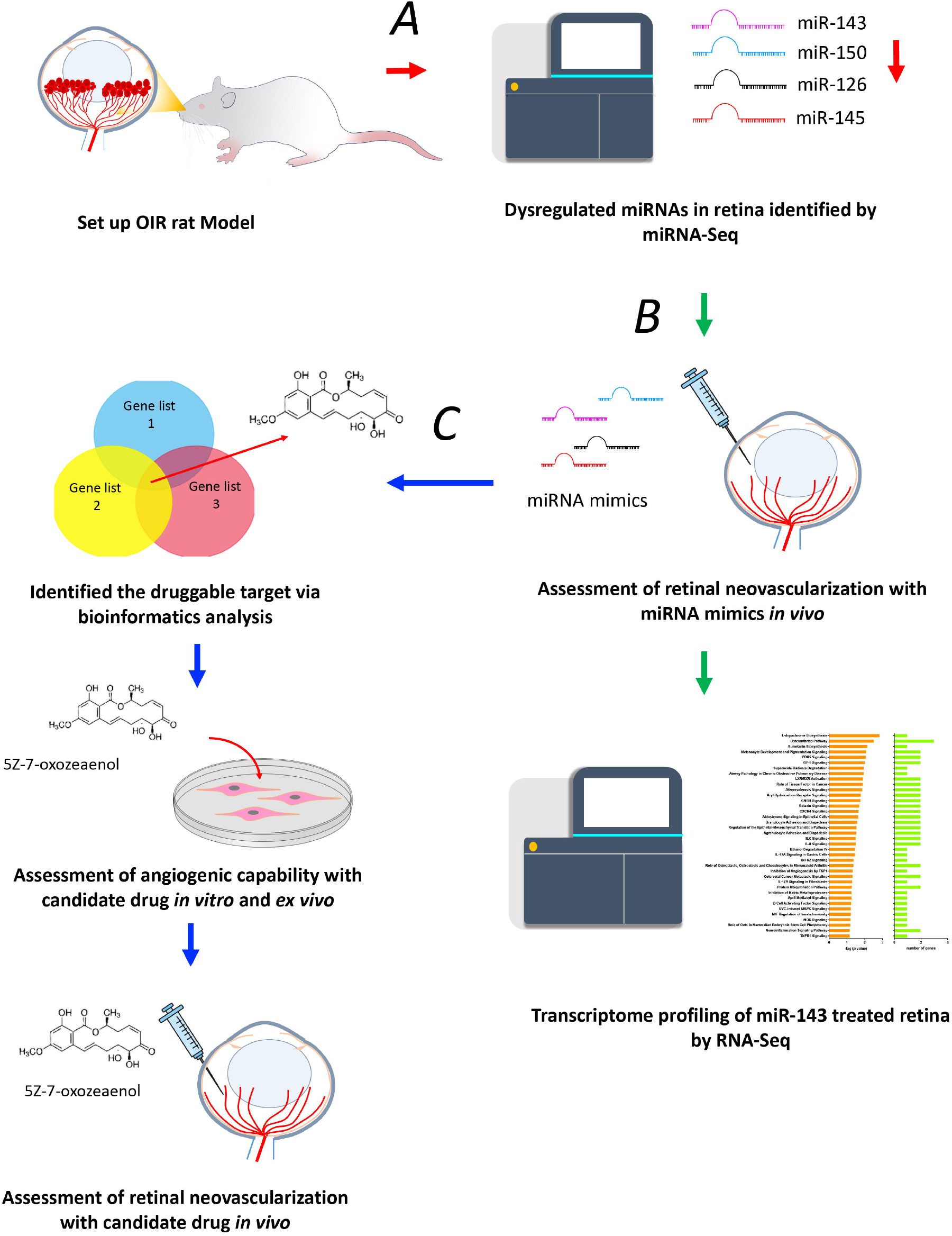
Schematic diagram of the stepwise miRNA identification protocol that informed the following *in vivo* therapeutic approach. (*A*) To identify the miRNAs associated with retinal neovascularization, we used a rat model of OIR and extracted the retinal miRNAs at P14 for miRNA-Seq. Four miRNAs including miR-143, miR-150, miR-126, and miR-145 were down-regulated in OIR rats compared to normoxic rats. *(B)* To examine if restoration of down-regulated miRNAs can attenuate retinal neovascularization, we injected miRNA mimics intravitreally and assessed the efficacy. Retinal RNA was extracted from OIR rats receiving miR-143 mimics and RNA-Seq was performed to profile gene expression. Several genes were significantly changed in OIR rats receiving miR-143 mimics compared to those receiving scrambled RNAs treatment. The interaction of these gene products showed that the major gene cluster involves in the suppression of retinal neovascularization by miR-143. *(B)* Analysis *in silico* identified TAK1 as a potential mediator of retinal neovascularization that may be regulated by miR-143. A pharmacological inhibitor of TAK1, 5Z-7-Oxozeaenol, was tested for its anti-angiogenic actions *in vitro, ex vivo* and *in vivo*.

## Materials and Methods

### Animals

All experimental studies were performed according to the ARVO Statement for the Use of Animals in Ophthalmic and Vision Research. This study was approved by the Animal Ethics Committees of St Vincent’s Hospital (Melbourne) and the University of Melbourne (Reference Number 004/16 and 13-044UM). Pregnant Sprague-Dawley (female, 12-14 weeks old) and Brown Norway (male, 12 weeks old) rats were supplied by the Animal Resources Centre, Perth, Australia, and housed at the Experimental Medical and Surgical Unit, St Vincent’s Hospital (East Melbourne, Victoria, Australia) and Melbourne Brain Centre (Parkville, Victoria, Australia). The animal housing conditions are described in *Supplementary information.*

### OIR rat model and vessel quantification

OIR was induced as previously described(43). Briefly, newborn Sprague-Dawley rats and their nursing mothers were housed in custom-built, humidified chambers within 12 hours of birth (postnatal day 0, P0) and exposed to daily cycles of 80% O_2_ for 21 hours and room air for 3 hours from P0 to 14. The pups were then returned to room air until P18. An oxygen controller (PROOX 110; BioSpherix, Parish, NY, USA) was used to monitor and control the oxygen level. The rats were sacrificed at P18, and their retinas were dissected and stained with isolectin B_4_ (5 g/mL, Alexa Fluor 488, catalogue no. I21411; Life Technologies Australia, Mulgrave, VIC, Australia). The neovascularization and vaso-obliteration in the rat retinas were quantified using Adobe Photoshop (CC 2017.1.1)(44). The quantification and analysis were performed by two blinded assessors (JHW and LT).

### Extraction of RNA and qPCR

Total RNA and miRNA were isolated from the dissected retinas using miRNeasy mini kits (catalogue no. 217084; Qiagen, Chadstone, VIC, Australia). Quantitative PCR was performed using a StepOne™ Real-Time PCR System (Applied Biosystems, Foster City, CA, USA). The methods used are further described in *Supplementary information.* Detailed information regarding the qPCR assays is provided in Supplementary Table 1.

### MicroRNA and RNA next-generation sequencing

The microRNA and RNA next-generation sequencing was performed following established protocols(45; 46). Additional information is provided in *Supplementary information.*

### Computational analysis

To explore the miRNA regulatory network, we analyzed the putative target genes of the dysregulated miRNAs (i.e., miR-150-5p, miR-126-3p, miR-143-3p and miR-145-5p) that were validated by qPCR using Ingenuity Pathways Analysis 2016 (IPA 2016, Ingenuity System, http://www.ingenuity.com) and the core analysis method tool. The putative target genes of the down-regulated miRNAs were obtained from multiple online databases, including miRNA.org and IPA. All putative target genes were merged and entered into IPA for the core analysis. IPA analysis identified the canonical pathways related to the target genes of the down-regulated miRNAs by Fisher’s exact test and adjusted P-values<0.05. To reduce the range of target genes, the target genes in each selected signaling pathway were matched to the Gene Ontology-derived gene set(47) of angiogenesis (http://amigo.geneontology.org/amigo/term/GO:0001525). Thus, the angiogenic genes shared by several pathways were identified. IPA and core analysis were also applied to analyze the regulatory network of the genes selected from the RNA-Seq. To explore the functional properties of these identified genes, the STRING search tool (http://www.string-db.org/) was used to create protein-protein interaction networks and reveal the interactivity of these gene products.

### Intravitreal injection

Intravitreal injections of miRNA mimics (GE Healthcare Dharmacon, Parramatta, NSW, Australia; Supplementary Table 2) or 5Z-7-Oxozeaenol (catalogue no. 3604/1; Tocris Bioscience, Bristol, UK) were performed under a surgical microscope similar to that previously described(48; 49). Details regarding the intravitreal injections are provided in *Supplementary information.*

### Cell culture and assays

Human umbilical vein endothelial cells (HUVECs) were purchased from Lonza (catalogue no. CC-2519; Lonza, Walkersville, MD, USA) and cultured in endothelial cell basal medium-2 (EBM-2) supplied with EGM™-2 BulletKit™ (catalogue no. CC-5035; Lonza). HEK293A cells were purchased from Invitrogen (catalog no. R70507; Life Technologies Australia) and cultured in Dulbecco’s modified Eagle’s medium (DMEM) (catalog no. 11965118; Life Technologies Australia) supplemented with 10% fetal calf serum (Sigma-Aldrich, St. Louis, MO, USA), 2 mM glutamine (catalog no. 2503008; Life Technologies Australia), and 50 U/mL penicillin-streptomycin (catalog no. 15070063; Life Technologies Australia). Both cell lines tested negative for mycoplasma using a MycoAlert™ Mycoplasma Detection Kit (catalogue no. LT07; Lonza) and cultured in a humidified 5% CO_2_ atmosphere at 37 °C. HUVECs and HEK293A cells were used for endothelial functional assays (i.e., tube formation, migration, and proliferation) and luciferase assays, respectively. The details of the transfection and assay conditions are described in *Supplementary information*.

### Electroretinography (ERG)

A low dose (18 ng) of 5Z-7-Oxozeaenol or vehicle was injected intravitreally into the eyes of male Brown Norway rats; 28 days following the injection, the rats underwent ERG. The details of the functional assessment are described in *Supplementary information*.

### Optical coherence tomography (OCT)

Following the ERG measurement, the rat eyes were imaged using spectral domain-OCT. The details of this procedure are described in *Supplementary information*.

### Aortic ring sprouting assay

Aortic ring assays were performed following an established protocol(50). The aortae were isolated from C57BL/6 mice, sectioned into 1-mm-long rings, and cultured in Matrigel. The rings were imaged daily, and the sprouting area was quantified. Further information is provided in *Supplementary information.*

### Data availability

The retinal miRNA profiles of the rats and mice and retinal miRNA transcriptomes from the rat model of OIR by next-generation sequencing have been deposited in NCBI’s Gene Expression Omnibus and are accessible through GEO Series accession number GSE104593 (https://www.ncbi.nlm.nih.gov/geo/query/acc.cgi?acc=GSE104593) and GSE104620 (https://www.ncbi.nlm.nih.gov/geo/query/acc.cgi?acc=GSE104620), respectively. The RNA-Seq data can be obtained from NCBI’s Gene Expression Omnibus (https://www.ncbi.nlm.nih.gov/geo/query/acc.cgi?acc=GSE104588). The data that support the findings of this study are available in the supplementary information files and from the corresponding author upon request.

### Statistical analysis

All statistical analyses were performed using Prism 6 software (GraphPad Software, Inc., La Jolla, CA, USA). The results are represented as the mean±s.e.m. unless noted otherwise. P-values<0.05 were considered statistically significant. The P-values are represented as follows: *P<0.05 and **P<0.01.

## Results

### Retinal microRNA expression is significantly down-regulated in the OIR rats

Given mouse and rat models of OIR are commonly used for studying retinal neovascularization, we sought to identify which of the two species gives a higher degree of homology in retinal miRNA expression profile to human. Using miRNA-sequencing (miRN-Seq) by NGS, we found that under control conditions, both rats and mice shared ∼63.0% of human retinal miRNAs (8) (Supplementary Fig. 1A). By aligning the average read count of retinal miRNAs of rat or mouse with top 100 miRNAs expressed in the human retina (8), these miRNAs were found to be highly conserved between rat and mouse (Supplementary Fig. 1B). Therefore, we chose the rat model for our subsequent studies because the development of retinal neovascularization in the rats with OIR better reflects human retinopathy of prematurity (9; 10).

The OIR rat model involves exposing rat pups to daily cycles of 80% O_2_ for 21 hours and room air for 3 hours from postnatal day (P) 0 to P14. At P14, the rat pups are returned to a relatively hypoxic environment (room air), which results in pathologic neovascularization at P18 and regression of neovascularization by P29 (Fig. 2A). MicroRNA-Seq was performed to obtain an overall picture of the miRNA expression profile in the retinas of OIR rats and age-matched normoxic (housed in room air) rats at P14. A ranking of the identified miRNAs in the retina from normoxic rats based on the read count of each miRNA was listed in Supplemental Data 1. Seven miRNAs (miR-451-5p, miR-150-5p, miR-126-3p, miR-126-5p, miR-143-3p, miR-145-5p and miR-145-3p) were found to be down-regulated by the OIR with a Log2 (Fold Change) less than −1 and adjusted p value less than 0.05 compared to the normoxia (Table 1). Moreover, several miRNAs were upregulated but without statistical significance (adjusted p > 0.05) in the retinas from rats with OIR (Fig. 2B and 2C). Of those down-regulated miRNAs, we selected five miRNAs with mature miRNA strands for further analysis, which included miR-451-5p, miR-150-5p, miR-126-3p, miR-143-3p and miR-145-5p. Most importantly, these down-regulated miRNAs in the OIR rats at P14 were restored at P29 when the cyclic oxygen exposed neonatal rats were returned to room air for 15 days (Supplementary Table 3).

**Figure 2.**
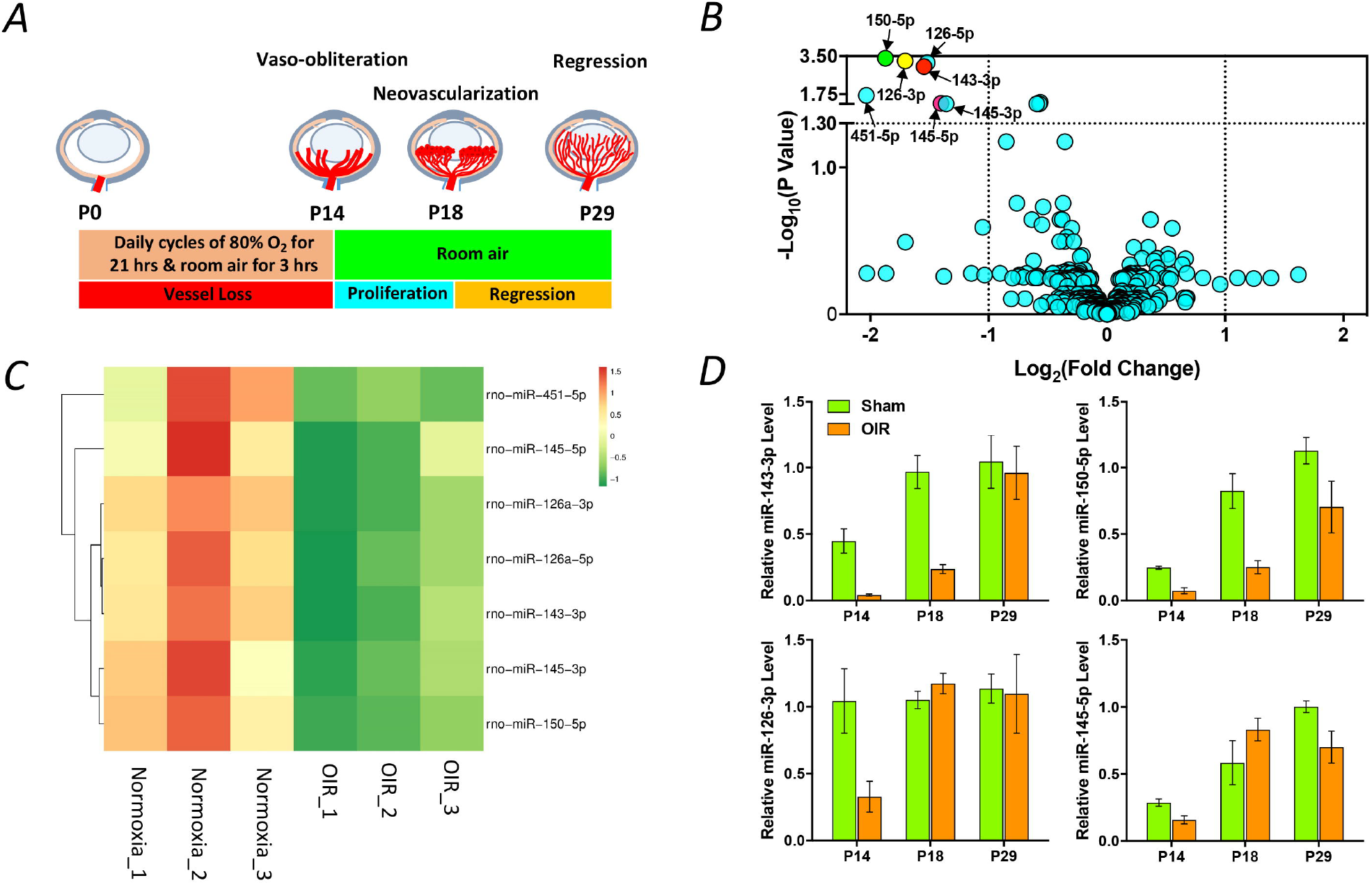
MicroRNAs are significantly down-regulated in the retinas of OIR rats. Schematic diagram of establishment of the OIR rat model. Neonatal rats were exposed to a daily cycle of 80% oxygen for 21 hours and room air for 3 hours from postnatal day (P) 0 to P14 and subsequently returned to room air. Retinal vaso-obliteration and neovascularization were observed in the OIR rats at P14 and P18, respectively. Retinal neovascularization was completely regressed, and the vasculature was fully normalized in the OIR rats at P29 (*A*). Total RNA, including miRNAs, was isolated from the retinas of OIR rats and age-matched normoxic rats at P14 and subsequently subjected to miRNA-Seq. The results are representative of three biological replicates (retinas) from one littermate. A volcano plot illustrates each miRNA with Log2 (Fold Change) (OIR versus normoxia) and p value (−Log10 (P-value) converted, e.g. −Log10 (0.05) = 1.3). Of 466 miRNAs, 7 were down-regulated with Log2 (Fold Change) less than −1 in the OIR rats compared to the normoxic rats. Of 7 miRNAs, miRNAs with mature form including miR-143-3p, miR-150-5p, miR-126-3p and miR-145-5p, were subjected to qPCR validation (*B*). A heat map illustrating the miRNAs that were significantly down-regulated in the retina of the OIR rats at P14 as analyzed by miRNA-Seq. Red indicates up-regulated expression levels; green indicates down-regulated expression levels.(*C*) Expression of retinal miR-150-5p, miR-126-3p, miR-143-3p, and miR-145-5p was suppressed in the OIR rats compared with those in the normoxic rats at P14 as analyzed by qPCR (*D*). All data were normalized to U6 small nuclear RNA. The results are representative of three biological replicates (retinas) from another littermate and expressed as the mean□±□s.e.m.

**Table 1.**
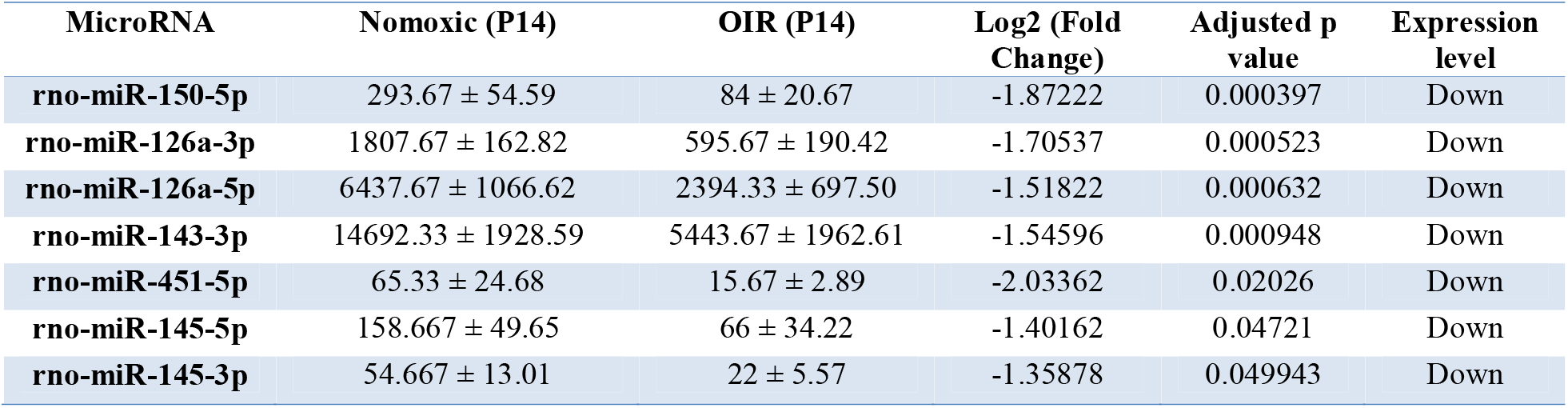
MicroRNAs are significantly down-regulated in the OIR rats at P14 compared to the normoxic rats.

| MicroRNA | Nomoxic (P14) | OIR (P14) | Log2 (Fold Change) | Adjusted p value | Expression level |
| --- | --- | --- | --- | --- | --- |
| **rno-miR-150-5p** | 293.67 ± 54.59 | 84 ± 20.67 | -1.87222 | 0.000397 | Down |
| **rno-miR-126a-3p** | 1807.67 ± 162.82 | 595.67 ± 190.42 | -1.70537 | 0.000523 | Down |
| **rno-miR-126a-5p** | 6437.67 ± 1066.62 | 2394.33 ± 697.50 | -1.51822 | 0.000632 | Down |
| **rno-miR-143-3p** | 14692.33 ± 1928.59 | 5443.67 ± 1962.61 | -1.54596 | 0.000948 | Down |
| **rno-miR-451-5p** | 65.33 ± 24.68 | 15.67 ± 2.89 | -2.03362 | 0.02026 | Down |
| **rno-miR-145-5p** | 158.667 ± 49.65 | 66 ± 34.22 | -1.40162 | 0.04721 | Down |
| **rno-miR-145-3p** | 54.667 ± 13.01 | 22 ± 5.57 | -1.35878 | 0.049943 | Down |

To validate the changes in the identified miRNAs, total retinal miRNA was isolated from an additional two cohorts of OIR and normoxic rats, and the miRNA expression was evaluated by performing quantitative PCR (qPCR). The expression profiles of miR-150-5p, miR-126-3p, miR-143-3p and miR-145-5p (Fig. 2D) were similar to those identified in the miRNA-Seq; however, the expression of miR-451-5p was lower and more variable in the OIR rats, and thus it was excluded from further analysis.

### MicroRNA treatment attenuates retinal neovascularization

To validate the involvement of the endogenous miRNA down-regulation in OIR rats, we quantified the effects of intravitreal injection of synthetic miR-143, miR-126, miR-150 and miR-145 mimics on retinal neovascularization at P14 (Fig. 3A). Four days after injection, retinal neovascularization was significantly attenuated in the OIR rats receiving a single intravitreal injection of synthetic miR-143, miR-126 or miR-150 mimics. However, no benefit was found in the OIR rats receiving miR-145 mimics (Fig. 3B, 3C and Supplementary Fig. 2). Moreover, no statistical difference was found in the vaso-obliteration (avascular area) between the rats receiving miRNA mimics and scramble RNAs in each group (Fig. 3D and Supplementary Fig. 2).

**Figure 3.**
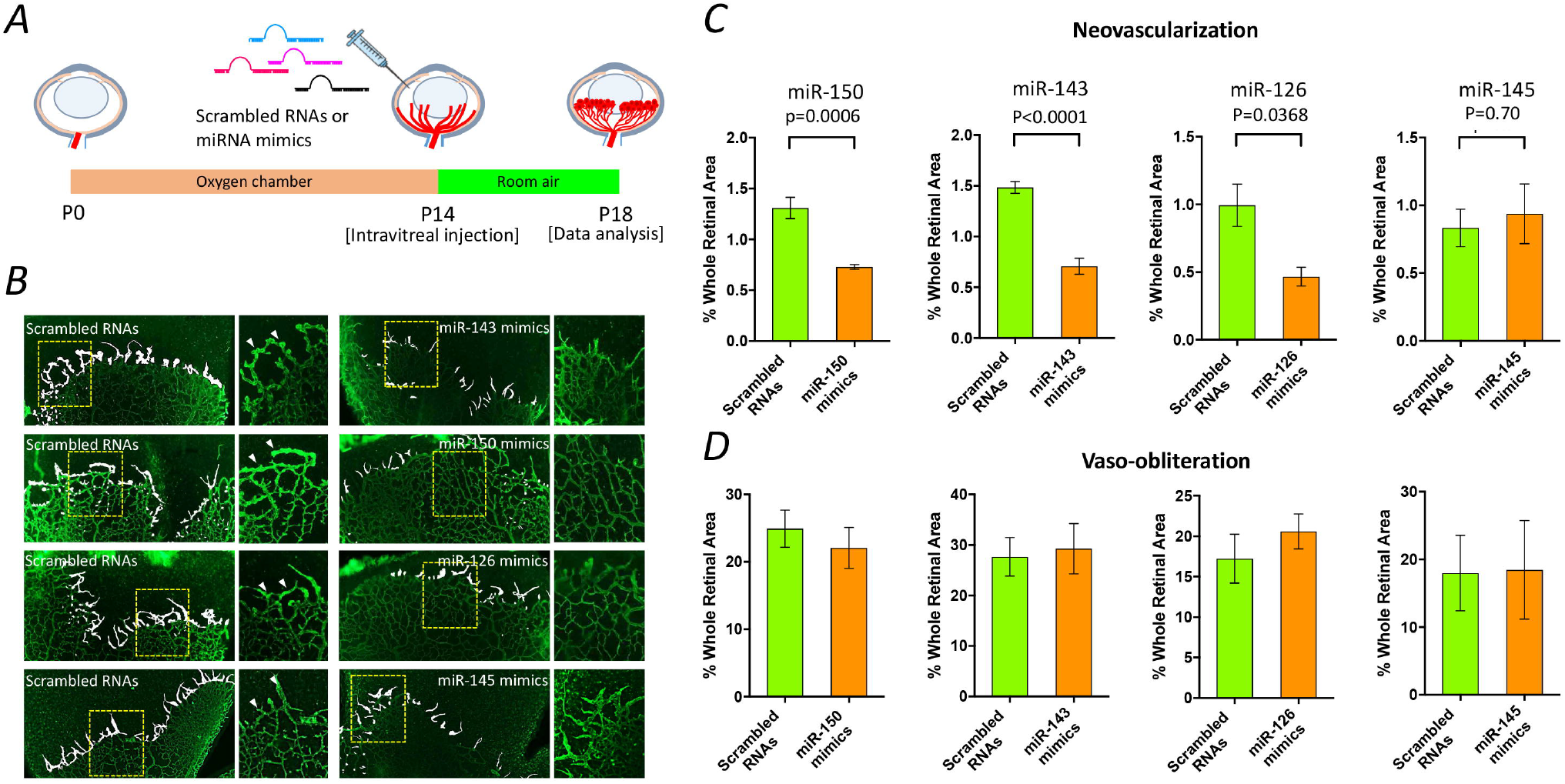
MicroRNA mimics alleviate retinal neovascularization *in vivo*. Schematic representation of miRNA mimic treatment in OIR rats. Scrambled RNAs or miRNA mimics (1 µg) was injected intravitreally into OIR rats at P14, and a quantitative analysis was performed at P18 (*A*). Representative images of retinal flat-mount 4 days after a single intravitreal injection of miRNA mimics. Retinal neovascularization is highlighted in white, and insets show the selected areas at a high magnification. The arrows indicate vessel tufts (*B*). Quantitative analysis of retinal neovascularization and vaso-obliteration in OIR rats at P18. The results are representative of five biological independent replicates (retinas) from two littermates per group. Two-tailed Student’s *t*-test was performed to determine the significance of the differences. No significant difference was observed in vaso-obliteration, two-tailed Student’s *t*-test: P=0.5075 for miR-150; P=0.8029 for miR-143; P=0.4148 for miR-126; P=0.9620 for miR-145. Data are expressed as the mean□±□s.e.m (*C* and *D*).

### MicroRNA-143 suppresses retinal neovascularization via inflammation/stress-related pathways

Since miR-143 is highly enriched in the normal human retina and the role of miR-143 has never been explored, we sought to explore the underlying mechanism by which miR-143 attenuates retinal neovascularization in OIR rats. Neonatal pups subjected to either normoxia or OIR for 14 days (P14) were received an intravitreal injection of either scrambled miRNAs (control) or miR-143 mimics, and the retinas of rats were dissected at P18. Total RNAs were isolated from the retinas and subjected to RNA-Seq (Fig. 4A). The following experimental design compared 4 groups of rats with different retinal vasculature and miRNA treatments: 1) Normoxic-control: normoxic rats receiving non-targeting scrambled RNAs; 2) Normoxic-miR-143: normoxic rats receiving miR-143 mimics; 3) OIR-control: OIR rats receiving non-targeting scrambled RNAs; 4) OIR-miR-143: OIR rats receiving miR-143 mimics.

**Figure 4.**
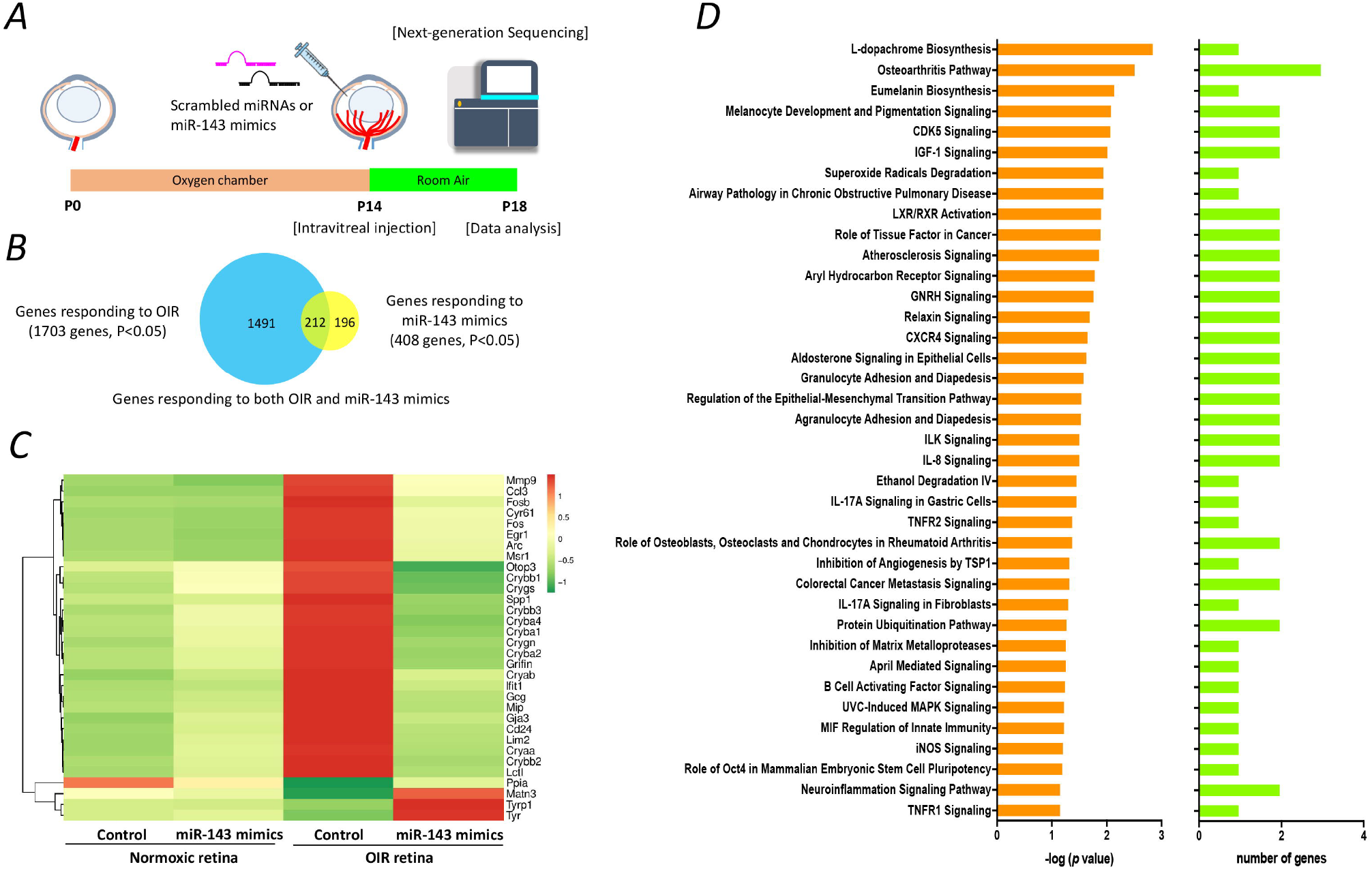
miR-143 indirectly regulates target genes and attenuates retinal neovascularization. Schematic representation of miR-143 mimic treatment in OIR rats (n=3 per group from one littermate). Non-targeting scrambled RNAs (control) or miR-143 mimic (1 µg) was injected intravitreally into the OIR or normoxic rats at P14, and RNA-Seq was performed at P18 (*A*). Computational analysis identified 1,703 genes differentially expressed in the retina between the OIR rats and normoxic rats (adjusted P<0.05), both of which receiving scrambled RNAs. Moreover, 408 genes (adjusted P<0.05) were identified with significantly altered expression in the OIR rats receiving miR-143 mimics compared with the OIR rats receiving scrambled RNAs. Venn diagram depicts the overlap between these two gene clusters. Over 200 genes were identified that presumably involve in the mechanism by which miR-143 mimics suppressed retinal neovascularization in OIR rats. (*B*). A heatmap illustrating the expression levels of selected genes responding to miR-143 mimics in the OIR rats. The mean expression levels of selected 32 genes were significantly altered in the retinas of OIR rats receiving scrambled RNAs at P18, whereas their expression level was reversed after receiving miR-143 mimics. Red indicates up-regulated expression levels; green indicates down-regulated expression levels. Control: scrambled miRNA (*C*). Protein-protein-interaction analysis by STRING suggested that two major protein clusters mediated by *Fos* and *Cryaa* may play critical roles in suppression of retinal neovascularization by miR-143 mimics treatment in the OIR rats (*D*).

Analysis of RNA-Seq data identified 1703 genes differentially expressed between the OIR-control and normoxic-control retinas (adjusted P<0.05, Supplemental Data 2) and 408 genes with altered levels between the OIR-miR-143 and OIR-control retinas (adjusted P<0.05, Supplemental Data 3). Of these genes, a total of 212 genes were found to be differentially expressed in both comparisons (Fig. 4B) and thus are likely to be important in retinal neovascularization and are regulated by miR-143. Of the 212 genes, 71 were differentially expressed in the OIR-control retinas, with their expression levels were normalized in the OIR rats receiving miR-143 mimics. Further filtering the genes based on significant change in expression (Log2(Fold Change) > +1.0 or < −1.0) narrowed down the list to 32 genes (Fig. 4C and Table 2). We also found that the expression levels of these genes were either reduced or increased in the OIR rats receiving miR-143 mimics compared to the OIR rats receiving scrambled RNAs. Notably, none of the identified genes was established as direct targets of miR-143 in the miRBase (http://www.mirbase.org/) or TargetScan (http://www.targetscan.org/vert_71/) databases, suggesting that these altered genes are potentially regulated by miR-143 in an indirect manner. The STRING database was used to identify known interactions between each of 32 genes. We identified two major gene clusters that could lead to a reduction of retinal neovascularization in the OIR rats receiving miR-143 mimics. The detailed gene product interactions are shown in Fig. 4D. Of 32 genes, Fos or αA-crystallin (Cryaa) was found to have the largest number of interactions. Pathway analysis suggested that the inflammation/stress-related pathways, such as IGF-1 signaling, GNRH signaling and CXCR4 signaling (Supplementary Fig. 3).

**Table 2.**
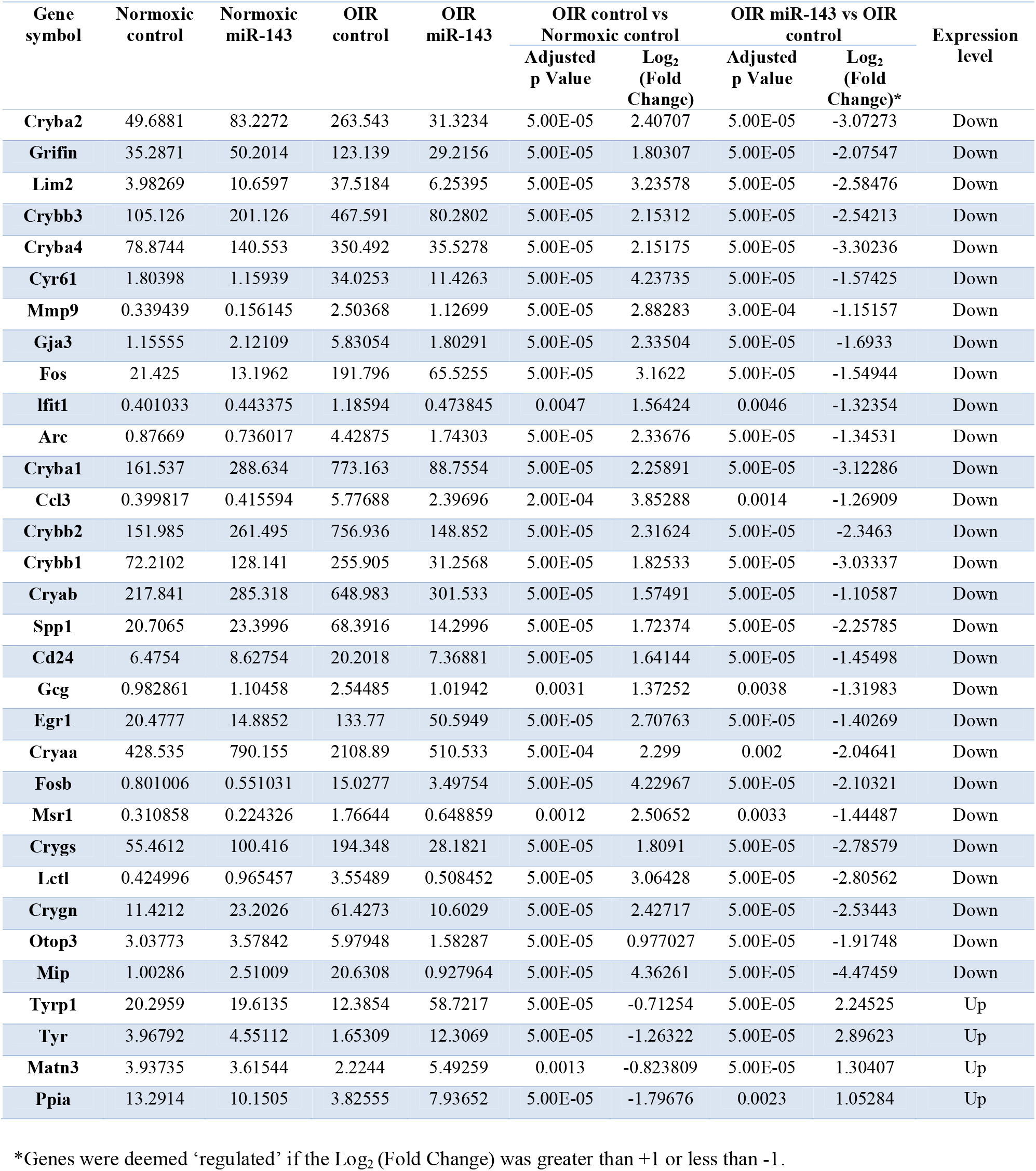
Expression of putative genes is significantly altered in the OIR rats receiving miR-143 mimics compared with those receiving non-targeting scrambled RNAs.

### Identification of *TAK1* as a potential therapeutic target in OIR

To interrogate the putative functions of the down-regulated miRNAs (miR-143-3p, miR-126-3p, miR-150-5p and miR-145-5p) in the OIR rats, implicated target genes of individual miRNA were predicted using online databases including miRNA.org and Ingenuity Pathways Analysis (IPA), and were subjected to a canonical pathway analysis. A total of 2,763 target genes involved in 38 different biological processes or cellular functions were identified (Fig. 5A). Three pathways associated with angiogenesis and diabetic retinopathy (11; 12) were identified, including molecular mechanisms of cancer and the transforming growth factor-beta (TGF-β) and AMP-activated protein kinase (AMPK) signaling pathways. To further refine the list of putative target genes, filters for the overlapping target genes involved in each signaling pathway and Gene Ontology-derived angiogenic gene set were applied. *TAK1* was identified as the sole pro-angiogenic gene involved in all three pathways (Fig. 5B). This gene is regulated by miR-143-3p. Other pro-angiogenic genes regulated by miR-143-3p, miR-126-3p, and miR-150-5p were also predicted (Supplementary Fig. 4).

**Figure 5.**
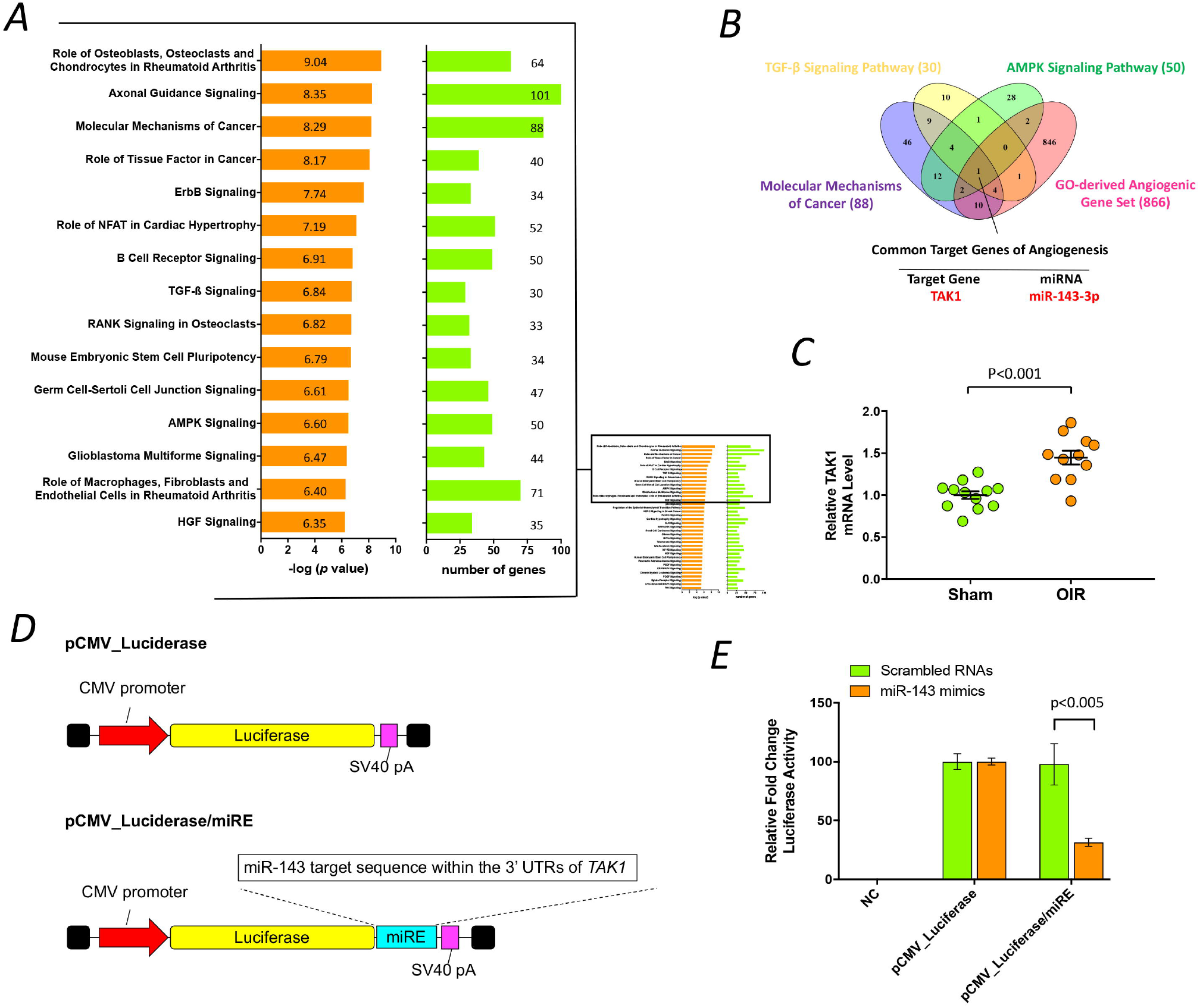
Prediction of target genes regulated by down-regulated miRNAs in OIR rats. Through *in silico* analysis by Ingenuity Pathway Analysis (IPA), a range of canonical pathways were identified and predicted to be relevant to down-regulated miRNAs, including miR-143-3p, miR-126-3p, miR-150-5p and miR-145-5p. These pathways emerged following IPA Core Analysis. Left column in the diagraph shows category scores; the threshold indicating the minimum significance level scored as −Log(p-value) from Fisher’s exact test, set here to 1.25. Right column in the diagraph refers to the number of molecules from the dataset that map to the canonical pathway from within the IPA knowledgebase. (*A*). Three pathways associated with retinal neovascularization, including molecular mechanisms of cancer and the TGF-β signaling and AMPK signaling pathways, were selected for further computational analysis. The Venn diagram depicts the overlap between predicted target genes involved in the three pathways and the Gene Ontology (GO)-derived angiogenic gene set. *TAK1*, a putative target gene of miR-143-3p, was identified as the sole pro-angiogenic gene involved in all three pathways. (*B*). TAK1 mRNA was significantly up-regulated in the retina of OIR rats at P14 compared with that in the normoxic rats as analyzed by qPCR. The results are representative of twelve and eleven independent biological replicates (retinas, three different littermates per group) from the normoxic and OIR groups, respectively. Data are expressed as the mean□±□s.e.m Two-tailed Student’s *t*-test was performed to determine the significant difference (*C*). The diagram of reporter vectors that encoding firefly luciferase with or without miR-143 responding elements from 3’-UTR of TAK1 gene (pCMV_Luciferase/miRE or pCMV_Luciferase). UTR: untranslated region. miRE: miRNA response elements. The expression levels of firefly luciferase were suppressed in HEK293A cells that co-transfected the reporter vector containing the target sequence of *TAK1* along with the miR-143 mimics. The results are representative of four independent biological replicates from two experiments and are expressed as the mean±s.e.m. Two-way ANOVA, followed by Tukey’s test, was performed to determine the significance of the differences (P=0.0006 for the interaction, P=0.0033 for treatment, P<0.0001 for plasmids; **P<0.005) (*D*).

A qPCR analysis was conducted to validate the retinal mRNA expression of *TAK1* in the retina of OIR rats. The *TAK1* mRNA expression was significantly increased in the OIR rats at P14 but normalized at P29 relative to the normoxic rats (Fig. 5C). The increased *TAK1* mRNA expression in the OIR rats at P14 was likely associated with the decreased expression of miR-143 as *TAK1* is negatively regulated by miR-143. To examine the regulatory effect of miR-143-3p on *TAK1* expression, we co-transfected a reporter vector encoding the firefly luciferase gene, followed by the miR-143-3p target sequence from the 3’ UTR of *TAK1* gene, along with a synthetic miR-143 mimic into HEK293A cells. The luciferase activity was markedly suppressed in the cells transfected with the reporter vectors containing the target sequences of *TAK1* (Fig. 5D, 5E and Supplementary Table 4). These results suggested that miR-143 directly regulates the expression of *TAK1*, at least in part, at the post-transcriptional level through targeting the 3’UTR target sequence of *TAK1 gene*.

### Pharmacological inhibition of TAK1 suppresses angiogenesis *in vitro* and *ex vivo* and retinal neovascularization *in vivo*

Subsequently, we sought to validate the effect of TAK1 inhibition on angiogenic processes *in vitro*. We investigated the effect of a commercially available TAK1 inhibitor, 5Z-7-Oxozeaenol, on angiogenesis in human umbilical vein endothelial cells (HUVECs). Compared with cells treated with the vehicle alone, cell proliferation was significantly inhibited (18.7% and 46.1% reduction at 24 and 48 hours, respectively) in the presence of 1000 nM 5Z-7-Oxozeaenol (Fig. 6A). The cells treated with 5Z-7-Oxozeaenol showed reduced migration in the scratch migration assay (63.2% reduction, Fig. 6B) compared with the vehicle-treated cells. Moreover, the cells treated with 1000 nM 5Z-7-Oxozeaenol showed a significant (56.2%) reduction in branching compared with the vehicle-treated cells (Fig. 6C). Nine days of treatment with 5Z-7-Oxozeaenol also suppressed vascular sprouting from aortic ring explants isolated from C57BL/6 mice (45.8% reduction, Fig. 6D). Altogether, these *in vitro* and *ex vivo* data indicate that the inhibition of TAK1 suppresses angiogenic activity.

**Figure 6.**
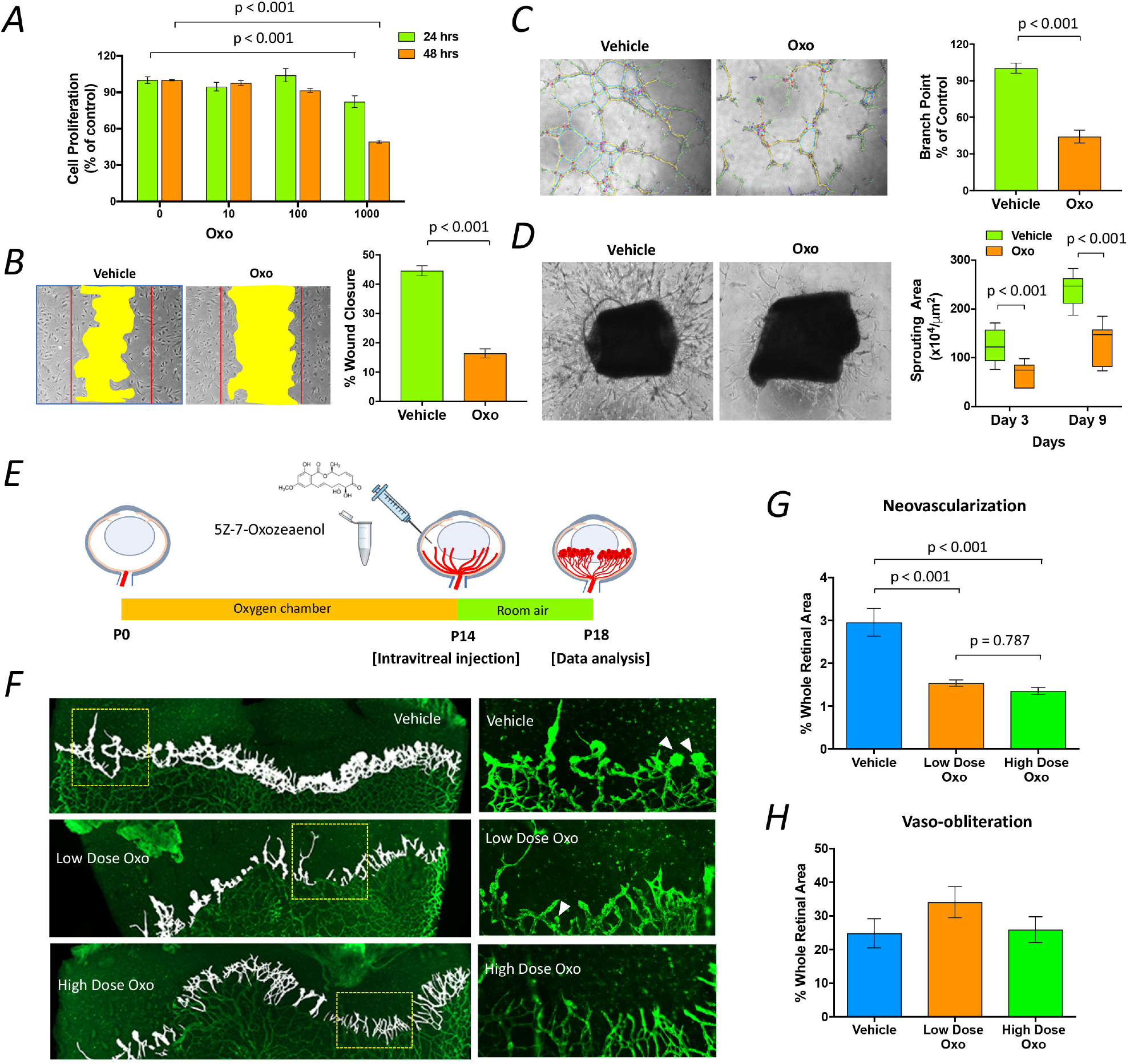
Inhibition of TAK1 by 5Z-7-Oxozeaenol suppresses angiogenesis *in vitro, ex vivo* and *in vivo*. HUVECs treated with 1000 nM 5Z-7-Oxozeaenol had a lower proliferative capability after 24 hours and 48 hours. Two-way ANOVA, followed by Tukey’s test, was performed to determine the significance of the differences (P<0.0001 for the interaction, P<0.0001 for concentration, P<0.0001 for time) (*A*). Representative images and quantitative analysis of cell migration assay characterizing wound closure. The results are representative of eight independent biological replicates from three experiments. Two-tailed Student’s *t*-test was performed to determine the significance of the differences (*B*). Representative images and quantitative analysis of tube formation assay characterizing the branch point numbers. The results are representative of twelve independent biological replicates from three experiments. Two-tailed Student’s *t*-test was performed to determine the significant difference (*C*). Representative images and quantitative analysis of vascular sprouting in 10-week-old C57BL/6 mice aortic ring explants. The results are representative of six independent biological replicates from two experiments. Two-way ANOVA, followed by Tukey’s test, was performed to determine the significance of the differences (P=0.0134 for the interaction, P=0.0010 for treatment, P<0.0001 for time) (*D*). Schematic representation of 5Z-7-Oxozeaenol treatment in OIR rats. Low (18 ng) or high (90 ng) dose of 5Z-7-Oxozeaenol was injected intravitreally into OIR rats at P14, and a quantitative analysis was performed at P18 (*E*). Representative images of retinal flat mount 4 days after a single intravitreal injection of 5Z-7-Oxozeaenol. Retinal neovascularization is highlighted in white, and insets show the selected areas at a high magnification. The arrows indicate vessel tufts (*F*). Quantitative analysis of retinal neovascularization and vaso-obliteration in OIR rats at P18. The results are representative of ten independent biological replicates (retinas) from three littermates per group. One-way ANOVA, followed by Tukey’s test, was performed to determine the significance of the differences. No significant difference was observed in vaso-obliteration. One-way ANOVA, followed by Tukey’s test, was performed to determine the significance of the differences (P=0.2658 for treatment). Data are expressed as the mean±s.e.m. Oxo: 5Z-7-Oxozeaenol (*G* and *H*).

To further examine the effect of TAK1 inhibition on retinal neovascularization, 5Z-7-Oxozeaenol was administered to the OIR rats via intravitreal injection at P14. The extent of neovascularization was quantified at P18 (Fig. 6E). A significant reduction in retinal neovascularization was observed in the OIR rats receiving either a low (18 ng/eye) or high (90 ng/eye) dose of 5Z-7-Oxozeaenol (50.0% and 53.3% reduction, respectively) compared with those receiving vehicle alone (Fig. 6F, 6G and Supplementary Fig. 5). No significant difference in vaso-obliteration was found among the three groups (Fig. 6H). We also evaluated whether 5Z-7-Oxozeaenol is toxic to the rodent retinas. Electroretinography (ERG) and optical coherence tomography (OCT) were performed to examine retinal functioning and structure 28 days after intravitreal injections of low-dose 5Z-7-Oxozeaenol in pigmented Brown Norway rats. No significant difference was observed in retinal function (Fig. 7A-D) or thickness (Fig. 7E-I) between the 5Z-7-Oxozeaenol and vehicle-treated eyes.

**Figure 7.**
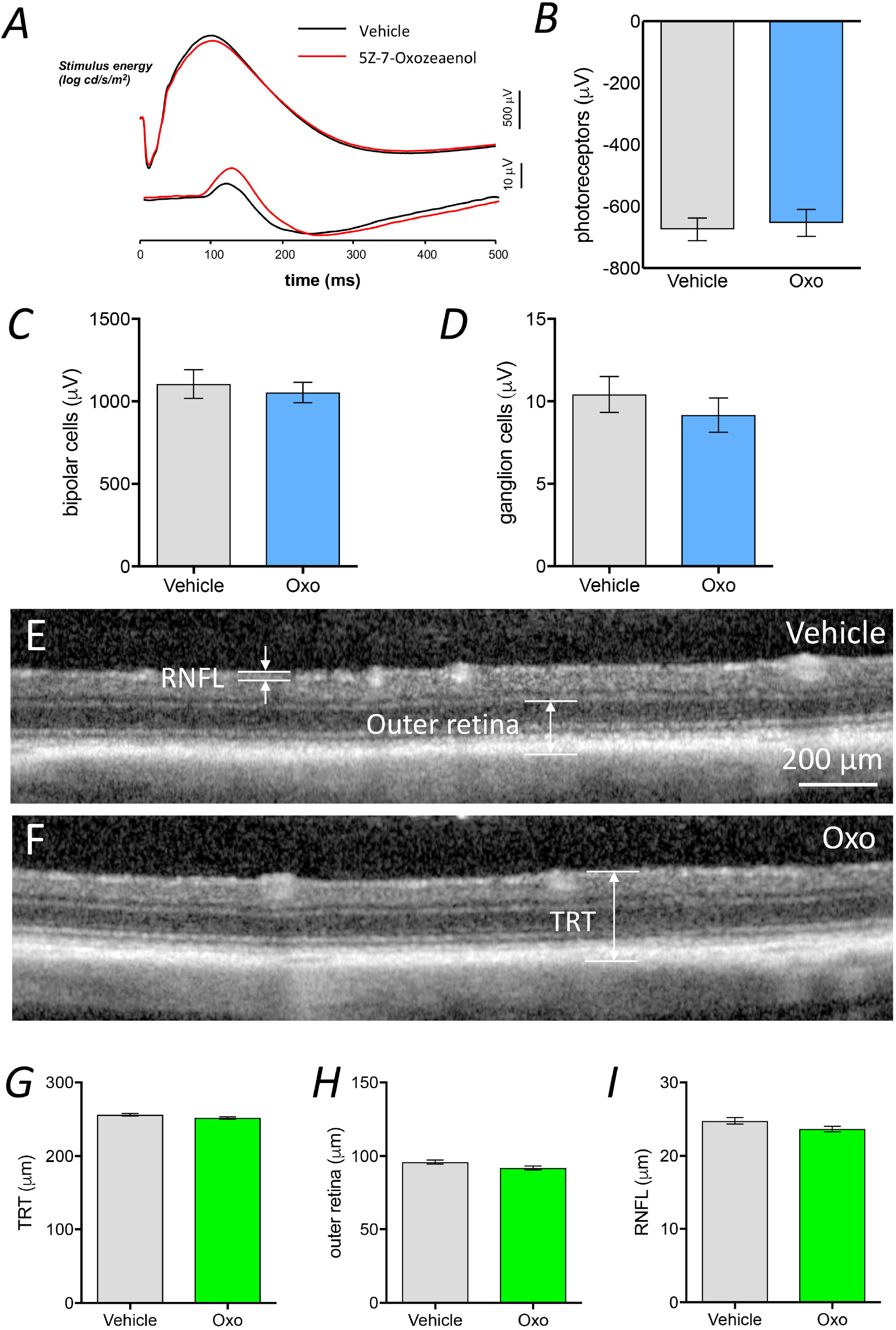
Effect of 5Z-7-Oxozeaenol on *in vivo* retinal structure and function. ERG and OCT were performed 28 days after intravitreal injection of 5Z-7-Oxozeaenol in Brown Norway rats. No difference was observed between the vehicle-(black traces) and 5Z-7-oxozeaenol-injected (red traces) eyes in terms of the average dim and bright flash ERG responses (*A*). Average photoreceptoral response amplitude of 5Z-7-Oxozeaenol-injected eyes and corresponding vehicle-injected eyes (the results are representative of 12 eyes, two-tailed Student’s *t*-test, P=0.4413) (*B*). Average bipolar cell response amplitude of 5Z-7-Oxozeaenol-injected eyes and corresponding vehicle-injected eyes (the results are representative of 12 eyes, two-tailed Student’s *t*-test, P=0.4980) (*C*). Average ganglion cell response amplitude of 5Z-7-Oxozeaenol-injected eyes and corresponding vehicle-injected eyes (the results are representative of 12 eyes, two-tailed Student’s *t*-test, P=0.1751) (*D*). Representative OCT images of 5Z-7-Oxozeaenol- and vehicle-injected eyes. Total retinal thickness (TRT), outer retina and retinal nerve fiber layer (RNFL) thickness were measured on both the temporal and nasal sides of the nerve, and the average was calculated (*E* and *F*). Average total retinal thickness (representative data are shown for 12 eyes, two-tailed Student’s *t*-test, P=0.0639) (*G*). Average outer-retinal thickness (representative data are shown for 12 eyes, two-tailed Student’s *t*-test, P=0.0502) (*H*). Average retinal nerve fiber layer thickness (the results are representative of 12 eyes, two-tailed Student’s *t*-test, P=0.0640). Data are expressed as the mean±s.e.m. Oxo: 5Z-7-Oxozeaenol (*I*).

## Discussion

MicroRNAs have been implicated in several *in vitro* and *in vivo* studies as crucial regulators of organ development and homeostasis (13). Many studies conducted over the previous decade suggest that abnormalities in miRNAs are involved in a range of disease processes, such as retinal neovascularization (14). Using miRNA-Seq, we identified four miRNAs including miR-143-3p, miR-126-3p, miR-150-5p and miR-145-5p were down-regulated in the retinas of rats subjected to ischemia-induced retinal angiogenesis. Of these down-regulated miRNA, miR-143-3p, miR-126-3p and miR-150-5p were directly contributed to the development of retinal neovascularization in OIR rats. Notably, our study provided the first evidence that dysregulation of miR-143 is involved in pathological retinal angiogenesis, and we also revealed that miR-143 alleviates retinal neovascularization by modulating the inflammation/stress pathway via *Fos* by RNA-Seq. Moreover, *in silico* analyses of miRNA-Seq data identified that TAK1 as a novel mediator involved in ischemia-induced retinal angiogenesis. The selective inhibition of TAK1 using 5Z-7-Oxozeaenol suppressed retinal neovascularization. Our data suggest that dysregulation of miRNAs is involved in the pathogenesis of retinal angiogenesis. Targeting the dysregulated miRNAs or inhibiting *TAK1* may be a feasible adjunct therapeutic approach in patients with retinal neovascularization.

Profiling miRNA expression in human specimens is an emerging approach to characterize diseases and identify novel biomarkers or therapeutic targets for diagnosis and treatment (15; 16). Unlike those tissues that are relatively accessible for biopsy, human retina is difficult to sample for analysis. Therefore, animal models are commonly used to investigate the roles of miRNAs in retinal neovascularization. The rodent OIR model is widely adopted to study the pathogenesis of ischemia-induced retinal angiogenesis or to test novel therapies for retinal neovascularization. Recently, a few studies investigated retinal miRNA expression in OIR mice using microarray-based platform and showed changes in retinal miRNA expression that being associated with retinal neovascularization (17-19). miRNA-Seq can provide absolute miRNA quantification with high accuracy and sensitivity compared with microarray assays (20; 21). Using miRNA-Seq, we have shown that the miRNA expression profile in the rat retina is comparable to that in the mouse retina. In addition, we identified that several miRNAs were differentially expressed following oxygen exposure in the retinas of OIR rats.

Previous miRNA profiling studies have identified several down-regulated miRNAs in the retinas of OIR mice, including miR-150, miR-375, miR-129-5p, and miR-129-3p (17; 18). In the present study, miR-143-3p, miR-126-3p, miR-150-5p and miR-145-5p were down-regulated in the OIR rats at P14 compared with those in the normoxic rats. This discrepancy may reflect differences in the tissue-specific expression of miRNAs across species (22) as well as differences in the oxygen exposure protocol. Studies have shown that restoration of endothelial-specific miR-126 or miR-150 via its respective mimics in retina can significantly attenuate retinal neovascularization in OIR mice through directly targeting pro-angiogenic genes (such as *VEGF, IGF-2*, and *HIF-1*α for miR-126; *CXCR4, DLL4*, and *FZD4* for miR-150) (23; 24). In line with previous studies, our study demonstrated that restoration of miR-126 and miR-150 suppressed retinal neovascularization not only in OIR mice but also in OIR rats. Most significantly, similar reduction of retinal neovascularization was also in OIR rats receiving miR-143 mimics. A study indicated that miR-143 may act as a switch signaling vascular smooth muscle cells to undergo phenotypic modulation, resulting in dedifferentiation and an increased rate of cellular proliferation and migration (25). This transition is an important component of pathogenesis in cardiovascular diseases, and the expression of miR-143/145 directly correlates with human and animal models of cardiovascular diseases (25). In a recent study, miR-143/145 acted as a communication molecule between smooth muscle cells and endothelial cells and modulated the angiogenic and vessel stabilization properties of the endothelial cells, a process induced by TGF-β (26). It has been reported that miR-143 is down-regulated in diseases associated with angiogenesis particularly in cancers, such as colon and lung cancers (27). Indeed, a study found that over-expression of miR-143 decreased tumor growth and angiogenesis, putatively by targeting and inhibiting the *NRAS*, which is a RAS family oncogene (28). The profound reduction of retinal neovascularization in the OIR rats receiving miR-143 mimics led us to hypothesize that miR-143 could be similar to miR-126 and miR-150 that being largely reported, play crucial roles in retinal neovascularization through dysregulating its target gene expression. Notably, miR-143 is highly enriched in the normal human retinas (29), and we have demonstrated that miR-143 is highly expressed compared with the other down-regulated miRNAs, such as miR-126 and miR-150, in normoxic rat retinas (miR-143 ranks 33^rd^ while miR-126 and miR-150 rank 87^th^ and 173^rd^, respectively in retinal miRNAs of rats. Supplementary Data 1). Thus, targeting miR-143 may be an alternative option for suppression of retinal neovascularization other than targeting vascular specific miR-126 and miR-150.

Intravitreal injection of miR-143 mimics may normalize aberrant expression of *Fos*- and *Cryaa*-mediated genes, and thus result in the suppression of retinal neovascularization in OIR rats. *Cryaa* is a lens-abundant gene (30) and plays important roles in maintaining lens transparency (31). Therefore, we presumed that *Fos*-mediated genes are predominantly responding to miR-143 regulation, thus resulting in the suppression of retinal neovascularization. In addition, *Fos* is commonly shared amongst the most of the signaling pathways identified by IPA Core Analysis, suggesting that *Fos* might be a key gene to take part in retinal neovascularization. Interestingly, none of the Fos-mediated genes are direct targets of miR-143, implicating that miR-143 may regulate these genes via other mediators. Indeed, several studies have suggested that “secondary” miRNAs (miRNAs regulated by other miRNAs through transcription factors) contribute to modulation of mRNAs, particularly the “indirect” mRNA targets (32-34). A study showed that approximately 30-50% of the indirect cardiac mRNA target regulation was attributable to higher-order effects of “secondary” miRNAs regulated by miR-378 and miR-499 (32). Another study using mouse model of cardiac pathology demonstrated that the efficacy of antimiR-34 (expression of miR-34 is elevated in settings of heart diseases) is likely attributed to its regulation of secondary miRNAs, producing direct and indirect effects on target mRNAs (35). These effects also likely underlie the actions of the miR-143 mimic, resulting in the attenuation of retinal neovascularization in OIR rats.

*TAK1* was identified by computational analysis with those pooled genes associated with down-regulated miRNAs in this study. TAK1, a serine/threonine kinase, which is a member of the mitogen-activated protein kinase (MAPK) family, is activated by inflammation and is a direct target gene of miR-143 (36; 37). TAK1 is a critical mediator of numerous important cellular pathways, such as the TGF-β and AMPK pathways (38; 39), and has been identified as a key regulator of angiogenesis. Activators of TAK1, such as Toll-like receptor ligands and inflammatory cytokines, have been suggested as the major promoters of angiogenesis due to their regulation of endothelial cell proliferation and migration in adult animals (40). A study using mice with an endothelial cell-specific gene deletion of *TAK1* found that endothelial *TAK1* modulates embryonic angiogenesis by affecting endothelial cell survival and migration (41). A further *in vitro* study demonstrated that the down-regulation of *TAK1* expression by siRNA could attenuate endothelial cell proliferation, migration, and tube formation as well as vascular sprouting capability (42). Furthermore, this study also showed that TAK1 acted as an AMP-activated protein kinase α1 (AMPKα1) kinase and positively regulated angiogenesis by increasing the expression of the antioxidative mitochondrial enzyme, superoxide dismutase 2 (SOD2) (42). Likewise, another study showed the similar results that TAK1-deficient HUVECs exhibited reduced tube formation and less cell migration (41). Consistent with these previous findings, our *in vitro* and *ex vivo* results demonstrated that the selective inhibition of TAK1 activity by 5Z-7-Oxozeaenol suppressed the *in vitro* angiogenic activity of primary endothelial cells and retinal neovascularization *in vivo*. Besides, we did not find any apparent changes of retinal function and structure after intravitreal injection of 5Z-7-Oxozeaenol, suggesting that 5Z-7-Oxozeaeno is safe at least in rats. Thus, TAK1 may represent a suitable target for the development of new therapeutics for retinal neovascularization.

In summary, this study demonstrates that miR-150-5p, miR-126-3p, miR-143-3p, and miR-145-5p are down-regulated in the retina of rats with proliferative retinopathy. Furthermore, we show that the retina-enriched miR-143 plays a protective role in retinal neovascularization via *Fos*. Selectively antagonizing the activity of TAK1 by 5Z-7-Oxozeaenol suppresses pro-angiogenic pathways and reduces retinal neovascularization. These findings highlight the utility of computational analyses using NGS data in screening for novel genes involved in complex pathological processes, such as retinal neovascularization. Moreover, the identification of novel genes allows for the identification of therapeutic targets, potential pharmacological agents, and new therapeutic approaches.

## Acknowledgements

The authors declare no conflict of interest. This work was supported by grants from The National Health and Medical Research Council of Australia (NHMRC#1061912 and 1123329), The Ophthalmic Research Institute of Australia and The Rebecca L Cooper Medical Research Foundation. J.H.W. received a R.B. McComas Research Scholarship in Ophthalmology, Gordon P Castles Scholarship and a Melbourne Research Scholarship. P.v.W. received a University of Melbourne, Annemarie Mankiewicz-Zelkin Fellowship and a Clinical Investigator Award from the Sylvia & Charles Viertel Charitable Foundation. B.V.B. received an Australian Research Council Future Fellowship (FT130100388). G.J.D. received a NHMRC Principal Research Fellowship (#1003113). A.W.H. received a NHMRC Practitioner Fellowship (#1103329). The Centre for Eye Research Australia receives Operational Infrastructure Support from the Victorian Government.

## Author contributions

Conceptualization, J.H.W. and G.S.L. Methodology, J.H.W., D.L., L.T. and G.S.L. Formal Analysis, J.H.W., V.S., B.V.B., A.W.H and G.S.L. Investigation, J.H.W., D.L., L.T., F.L., S.M.P., Z.H., B.V.B. and G.S.L. Resources, M.R. Data Curation, J.H.W. and G.S.L. Writing-Original Draft, J.H.W., D.L. and G.S.L. Writing-Review & Editing, B.V.B., A.W.H., G.J.D., and P.V.W. Visualization, J.H.W. and G.S.L. Supervision, G.J.D., P.V.W. and G.S.L. Project Administration, J.H.W. and G.S.L. Funding Acquisition, G.J.D. and G.S.L.

## References

1. Campochiaro PA: Retinal and choroidal neovascularization. J Cell Physiol 2000;184:301–310

2. Ross EL, Hutton DW, Stein JD, et al.: Cost-effectiveness of aflibercept, bevacizumab, and ranibizumab for diabetic macular edema treatment: Analysis from the diabetic retinopathy clinical research network comparative effectiveness trial. JAMA Ophthalmology 2016;

3. McClelland AD, Kantharidis P: microRNA in the development of diabetic complications. Clinical Science 2014;126:95–110

4. Raghunath A, Perumal E: Micro-RNAs and their roles in eye disorders. Ophthalmic Research 2015;53:169–186

5. Liu CH, Sun Y, Li J, Gong Y, Tian KT, Evans LP, Morss PC, Fredrick TW, Saba NJ, Chen J: Endothelial microRNA-150 is an intrinsic suppressor of pathologic ocular neovascularization. P Natl Acad Sci USA 2015;112:12163–12168

6. Bai Y, Bai X, Wang Z, Zhang X, Ruan C, Miao J: MicroRNA-126 inhibits ischemia-induced retinal neovascularization via regulating angiogenic growth factors. Experimental & Molecular Pathology 2011;91:471–477

7. Yan L, Lee S, Lazzaro DR, Aranda J, Grant MB, Chaqour B: Single and Compound Knock-outs of MicroRNA (miRNA)-155 and Its Angiogenic Gene Target CCN1 in Mice Alter Vascular and Neovascular Growth in the Retina via Resident Microglia. Journal of Biological Chemistry 2015;290:23264–23281

8. Karali M, Persico M, Mutarelli M, Carissimo A, Pizzo M, Singh Marwah V, Ambrosio C, Pinelli M, Carrella D, Ferrari S, Ponzin D, Nigro V, di Bernardo D, Banfi S: High-resolution analysis of the human retina miRNome reveals isomiR variations and novel microRNAs. Nucleic acids research 2016;44:1525–1540

9. Gammons MV, Bates DO: Models of Oxygen Induced Retinopathy in Rodents. Methods in molecular biology 2016;1430:317–332

10. Barnett JM, McCollum GW, Fowler JA, Duan JJ, Kay JD, Liu RQ, Bingaman DP, Penn JS: Pharmacologic and genetic manipulation of MMP-2 and -9 affects retinal neovascularization in rodent models of OIR. Invest Ophthalmol Vis Sci 2007;48:907–915

11. Gerhardinger C, Dagher Z, Sebastiani P, Park YS, Lorenzi M: The transforming growth factor-beta pathway is a common target of drugs that prevent experimental diabetic retinopathy. Diabetes 2009;58:1659–1667

12. Kubota S, Ozawa Y, Kurihara T, Sasaki M, Yuki K, Miyake S, Noda K, Ishida S, Tsubota K: Roles of AMP-activated protein kinase in diabetes-induced retinal inflammation. Invest Ophthalmol Vis Sci 2011;52:9142–9148

13. Vidigal JA, Ventura A: The biological functions of miRNAs: lessons from in vivo studies. Trends in cell biology 2015;25:137–147

14. Gulyaeva LF, Kushlinskiy NE: Regulatory mechanisms of microRNA expression. Journal of translational medicine 2016;14:143

15. van de Bunt M, Gaulton KJ, Parts L, Moran I, Johnson PR, Lindgren CM, Ferrer J, Gloyn AL, McCarthy MI: The miRNA profile of human pancreatic islets and beta-cells and relationship to type 2 diabetes pathogenesis. PloS one 2013;8:e55272

16. Lu J, Getz G, Miska EA, Alvarez-Saavedra E, Lamb J, Peck D, Sweet-Cordero A, Ebert BL, Mak RH, Ferrando AA, Downing JR, Jacks T, Horvitz HR, Golub TR: MicroRNA expression profiles classify human cancers. Nature 2005;435:834–838

17. Shen J, Yang X, Xie B, Chen Y, Swaim M, Hackett SF, Campochiaro PA: MicroRNAs regulate ocular neovascularization. Molecular therapy : the journal of the American Society of Gene Therapy 2008;16:1208–1216

18. Liu CH, Wang Z, Sun Y, SanGiovanni JP, Chen J: Retinal expression of small non-coding RNAs in a murine model of proliferative retinopathy. Scientific reports 2016;6:33947

19. Wang Y, Wu S, Yang Y, Peng F, Li Q, Tian P, Xiang E, Liang H, Wang B, Zhou X, Huang H, Zhou X: Differentially expressed miRNAs in oxygeninduced retinopathy newborn mouse models. Mol Med Rep 2017;15:146–152

20. Hurd PJ, Nelson CJ: Advantages of next-generation sequencing versus the microarray in epigenetic research. Brief Funct Genomic Proteomic 2009;8:174–183

21. Pritchard CC, Cheng HH, Tewari M: MicroRNA profiling: approaches and considerations. Nat Rev Genet 2012;13:358–369

22. Roux J, Gonzalez-Porta M, Robinson-Rechavi M: Comparative analysis of human and mouse expression data illuminates tissue-specific evolutionary patterns of miRNAs. Nucleic acids research 2012;40:5890–5900

23. Liu CH, Sun Y, Li J, Gong Y, Tian KT, Evans LP, Morss PC, Fredrick TW, Saba NJ, Chen J: Endothelial microRNA-150 is an intrinsic suppressor of pathologic ocular neovascularization. Proceedings of the National Academy of Sciences of the United States of America 2015;112:12163– 12168

24. Bai Y, Bai X, Wang Z, Zhang X, Ruan C, Miao J: MicroRNA-126 inhibits ischemia-induced retinal neovascularization via regulating angiogenic growth factors. Exp Mol Pathol 2011;91:471– 477

25. Rangrez AY, Massy ZA, Metzinger-Le Meuth V, Metzinger L: miR-143 and miR-145: molecular keys to switch the phenotype of vascular smooth muscle cells. Circulation Cardiovascular genetics 2011;4:197–205

26. Climent M, Quintavalle M, Miragoli M, Chen J, Condorelli G, Elia L: TGFbeta Triggers miR-143/145 Transfer From Smooth Muscle Cells to Endothelial Cells, Thereby Modulating Vessel Stabilization. Circulation research 2015;116:1753–1764

27. Calin GA, Croce CM: MicroRNA signatures in human cancers. Nat Rev Cancer 2006;6:857–866

28. Wang L, Shi ZM, Jiang CF, Liu X, Chen QD, Qian X, Li DM, Ge X, Wang XF, Liu LZ, You YP, Liu N, Jiang BH: MiR-143 acts as a tumor suppressor by targeting N-RAS and enhances temozolomide-induced apoptosis in glioma. Oncotarget 2014;5:5416–5427

29. Ludwig N, Leidinger P, Becker K, Backes C, Fehlmann T, Pallasch C, Rheinheimer S, Meder B, Stahler C, Meese E, Keller A: Distribution of miRNA expression across human tissues. Nucleic acids research 2016;44:3865–3877

30. Kannan R, Sreekumar PG, Hinton DR: Novel roles for alpha-crystallins in retinal function and disease. Progress in retinal and eye research 2012;31:576–604

31. Santhoshkumar P, Karmakar S, Sharma KK: Structural and functional consequences of chaperone site deletion in alphaA-crystallin. Biochim Biophys Acta 2016;1864:1529–1538

32. Matkovich SJ, Hu Y, Dorn GW, 2nd: Regulation of cardiac microRNAs by cardiac microRNAs. Circ Res 2013;113:62–71

33. van Rooij E, Quiat D, Johnson BA, Sutherland LB, Qi X, Richardson JA, Kelm RJ, Jr., Olson EN: A family of microRNAs encoded by myosin genes governs myosin expression and muscle performance. Dev Cell 2009;17:662–673

34. van Rooij E, Sutherland LB, Qi X, Richardson JA, Hill J, Olson EN: Control of stress-dependent cardiac growth and gene expression by a microRNA. Science 2007;316:575–579

35. Ooi JYY, Bernardo BC, Singla S, Patterson NL, Lin RCY, McMullen JR: Identification of miR-34 regulatory networks in settings of disease and antimiR-therapy: Implications for treating cardiac pathology and other diseases. RNA biology 2017;14:500–513

36. Broglie P, Matsumoto K, Akira S, Brautigan DL, Ninomiya-Tsuji J: Transforming growth factor beta-activated kinase 1 (TAK1) kinase adaptor, TAK1-binding protein 2, plays dual roles in TAK1 signaling by recruiting both an activator and an inhibitor of TAK1 kinase in tumor necrosis factor signaling pathway. The Journal of biological chemistry 2010;285:2333–2339

37. Prakhar P, Holla S, Ghorpade DS, Gilleron M, Puzo G, Udupa V, Balaji KN: Ac2PIM-responsive miR-150 and miR-143 target receptor-interacting protein kinase 2 and transforming growth factor beta-activated kinase 1 to suppress NOD2-induced immunomodulators. The Journal of biological chemistry 2015;290:26576–26586

38. Heldin CH, Moustakas A: Signaling Receptors for TGF-beta Family Members. Cold Spring Harbor perspectives in biology 2016;8

39. Herrero-Martin G, Hoyer-Hansen M, Garcia-Garcia C, Fumarola C, Farkas T, Lopez-Rivas A, Jaattela M: TAK1 activates AMPK-dependent cytoprotective autophagy in TRAIL-treated epithelial cells. The EMBO journal 2009;28:677–685

40. West XZ, Malinin NL, Merkulova AA, Tischenko M, Kerr BA, Borden EC, Podrez EA, Salomon RG, Byzova TV: Oxidative stress induces angiogenesis by activating TLR2 with novel endogenous ligands. Nature 2010;467:972–976

41. Morioka S, Inagaki M, Komatsu Y, Mishina Y, Matsumoto K, Ninomiya-Tsuji J: TAK1 kinase signaling regulates embryonic angiogenesis by modulating endothelial cell survival and migration. Blood 2012;120:3846–3857

42. Zippel N, Malik RA, Fromel T, Popp R, Bess E, Strilic B, Wettschureck N, Fleming I, Fisslthaler B: Transforming growth factor-beta-activated kinase 1 regulates angiogenesis via AMP-activated protein kinase-alpha1 and redox balance in endothelial cells. Arterioscler Thromb Vasc Biol 2013;33:2792–2799

43. Deliyanti D, Miller AG, Tan G, Binger KJ, Samson AL, Wilkinson-Berka JL: Neovascularization is attenuated with aldosterone synthase inhibition in rats with retinopathy. Hypertension 2012;59:607–613

44. Connor KM, Krah NM, Dennison RJ, Aderman CM, Chen J, Guerin KI, Sapieha P, Stahl A, Willett KL, Smith LEH: Quantification of oxygen-induced retinopathy in the mouse: a model of vessel loss, vessel regrowth and pathological angiogenesis. Nat Protoc 2009;4:1565–1573

45. Nixon B, Stanger SJ, Mihalas BP, Reilly JN, Anderson AL, Dun MD, Tyagi S, Holt JE, McLaughlin EA: Next Generation Sequencing Analysis Reveals Segmental Patterns of microRNA Expression in Mouse Epididymal Epithelial Cells. PloS one 2015;10:e0135605

46. Trapnell C, Williams BA, Pertea G, Mortazavi A, Kwan G, van Baren MJ, Salzberg SL, Wold BJ, Pachter L: Transcript assembly and quantification by RNA-Seq reveals unannotated transcripts and isoform switching during cell differentiation. Nat Biotechnol 2010;28:511–515

47. Powell JA: GO2MSIG, an automated GO based multi-species gene set generator for gene set enrichment analysis. BMC bioinformatics 2014;15:146

48. Hewing NJ, Weskamp G, Vermaat J, Farage E, Glomski K, Swendeman S, Chan RV, Chiang MF, Khokha R, Anand-Apte B, Blobel CP: Intravitreal injection of TIMP3 or the EGFR inhibitor erlotinib offers protection from oxygen-induced retinopathy in mice. Invest Ophthalmol Vis Sci 2013;54:864–870

49. Chen J, Connor KM, Aderman CM, Willett KL, Aspegren OP, Smith LE: Suppression of retinal neovascularization by erythropoietin siRNA in a mouse model of proliferative retinopathy. Invest Ophthalmol Vis Sci 2009;50:1329–1335

50. Baker M, Robinson SD, Lechertier T, Barber PR, Tavora B, D’Amico G, Jones DT, Vojnovic B, Hodivala-Dilke K: Use of the mouse aortic ring assay to study angiogenesis. Nat Protoc 2012;7:89– 104

