## Supplemental Tables and Figures for "MicroRNA-143 plays a protective role in ischemia-induced retinal neovascularization"

Running title: Role of microRNA-143 in retinal neovascularization

**Supplemental Material- Additional detailed methods**

**Supplemental Material- Supplementary Figures**

**Supplementary Figure 1.** Rat and mouse have highly homologous miRNA expression in the retina.

**Supplementary Figure 2.** The uncropped images of the retinal flat-mount used in Figure 3B.

**Supplementary Figure 3.** Canonical pathways involved in the regulatory mechanism of miR-143 mimics treatment.

**Supplementary Figure 4.** Identification of pro-angiogenic target genes regulated by miR-143 or miR-150 or miR-126.

**Supplementary Figure 5.** The uncropped images of the retinal flat-mount used in Figure 6G.

**Supplemental Material- Supplementary Tables**

**Supplementary Table 1.** TaqMan probe sequences used for the real-time quantitative polymerase chain reaction.

**Supplementary Table 2.** Oligonucleotide sequences of miRNA mimic.

**Supplementary Table 3.** The expression levels of down-regulated miRNAs in OIR rats at P14 are restored at P29.

**Supplementary Table 4.** Oligonucleotide sequences for the luciferase reporter constructs.

**Supplementary Material– Additional detailed methods**

**Animal housing.** Animals were housed in standard cages, with free access to food and water under a 12 hours light (50 lux illumination) and 12 hour dark (<10 lux illumination) cycle with a temperature controlled environment to minimize possible light-induced damage to the eye.

**Extraction of total RNAs and microRNAs.** The eyes of OIR and normoxic rats were enucleated at P14 and P29 respectively and stored in the RNAlater (catalogue no. 76104; Qiagen, Chadstone, VIC, Australia), and the retinas were dissected immediately under a dissection microscope. Total RNAs and miRNAs were isolated from the dissected retina by using a miRNeasy mini kit (catalogue no. 217084; Qiagen). The quantity of RNA was measured by using a spectrophotometry, and the integrity of total RNA was confirmed by a Bioanalyzer at the Australian Genome Research Facility (AGRF, Melbourne, VIC, Australia). The quality of RNA was determined by RNA Integrity Number (RIN) and concentration of RNA. A RIN of > 7 and concentration of > 20 ng/µL are considered acceptable for miRNA sequencing. We paired qualified RNA samples (n=3) from OIR and normoxic rats at P14 for miRNA-Seq.

**MicroRNA next-generation sequencing.** One microgram of RNAs containing miRNAs in triplicate from the normoxic rats and OIR rats at P14 were prepared as per the manufacturers’ instructions (Illumina Inc. San Diego, CA, USA). The libraries of cDNA from the biological samples for normoxic and OIR groups were sequenced by using an Illumina Hiseq-2500 RNA-seq platform as 50 bp single end chemistry at Australian Genome Research Facility (AGRF, Melbourne, VIC, Australia) (1; 2). Briefly, the sequence reads from all samples were analysed for overall quality, screened for the presence of any contaminants and trimmed accordingly. The cleaned sequence reads were then processed through the quantification modules miRDEEP2 ver2.0.0.7 pipeline for known miRNA expression profiling. Rat miRNAs from miRBASE release21 were used for mapping and quantification (3). Differential miRNA expression analysis was undertaken using the eLimma bioconductor package (<https://www.bioconductor.org/packages/release/bioc/vignettes/limma/inst/doc/usersguide.pdf)>. Expression profiling comparisons were performed for mature miRNAs between the normoxic rats and OIR rats at P14 with a data filter set to ≤ -1 Log_2_ (Fold Change) and false discovery rate (FDR) of 0.05.

**RNA Next-generation Sequencing.** The libraries of cDNA preparation were similar to miRNA-Seq as above. The quality of the raw reads was assessed by using FASTQC (4) and we ensured that all reads had phred scores > 28. The reference genomic sequence of *Rattus norvegicus* (rn6) was retrieved from the iGenomes (obtained on 19^th^ June 2017, https://support.illumina.com/sequencing/sequencing_software/igenome.html) and used for all analysis. The quality reads were mapped to the reference genome using TopHat (5). After that, Cufflinks software was used to estimate gene expression level using Fragments per Kilobase of transcript per Million mapped reads (FPKM)(6). To identify genes, a Cuffdiff (6) statistical method was employed to calculate the differences of expression and transcript level between groups. A q-value was used to calculate differentially expressed genes between groups in the study. The tools above used in the analysis was hosted on Galaxy-Server (http://usegalaxy.org/). CummeRbund (7) transforms the Cuffdiff data into the R statistical computing environment, which was used for functional analysis and generating plots. Heatmaps were generated using a ClustVis (8). Functional analysis of genes and their networks were used in the analysis and the significance of any gene function in a network was denoted by an adjusted p-value of less than 0.05 (Fisher Exact Test). Genes were deemed to be ‘regulated’ if the Log2 (Fold Change) was greater than 1.0, or less than -1.0.

**Quantitative Polymerase Chain Reaction (qPCR)**

The miRNAs profiles and putative target genes selected as per the computational analysis were validated by qPCR with Taqman miRNA assay and gene assay reagents (Life Technologies Australia, Mulgrave, VIC, Australia) respectively according to the manufacturer’s instructions. Real-time quantitative PCR was performed using the StepOne™ Real-Time PCR Systems (Applied Biosystems, Foster City, CA, USA). Briefly, for miRNA reverse transcription (RT), 100 ng of total RNA was reversely transcribed to individual cDNA with individual miRNA RT primer using a miRNA RT kit (Life Technologies Australia). For mRNA reverse transcription, total RNA (100 ng) was reverse-transcribed to cDNA using a high-capacity RT kit (catalog no. 4374996; Life Technologies Australia). Quantitative PCR was then performed using a TaqMan faster PCR master mix and commercially available rat miRNA and gene probe sets (TaqMan Gene Expression Assay, Life Technologies Australia), respectively. The miRNAs selected for validation were miR-143-3p, miR-145-5p, miR-126-3p, miR-150-5p and miR-451-5p. The U6 small nuclear RNA was employed as an endogenous control to normalise the expression levels of miRNAs. The putative target genes selected for validation was TAK1. Rat beta-actin (ACTB) was used as a reference gene. For analysis of miRNA and gene expression, relative expression levels of miRNAs and target genes between normoxic and OIR rats were calculated using ΔΔCt method which has been previously described by Livak (9). All qPCR assays for validation of candidate miRNAs and genes were performed in duplicate using samples of cohorts that different from the one used for miRNA sequencing. The detailed information of qPCR assays is provided in Supplementary Table 1.

**Animal allocation.** Rats were equally randomly subjected to different treatment groups from same littermate or cage by investigators (JHW and GSL for OIR studies; SMP and BVB for ERG/OCT) and housing in same room at the animal facility.

**Intravitreal Injection.** Intravitreal injections were performed in normoxic rats and OIR rats at P14 using a surgical microscope similar to that previously described (10; 11). In brief, after making a guide track through the conjunctiva and sclera at the superior temporal hemisphere behind the limbus using a 30-gauge needle, a hand-pulled glass micropipette connected to a 10 μL Hamilton syringe (Bio-Strategy, Broadmeadows, VIC, Australia) was inserted into the vitreal cavity. One microliter of low (15 ng) or high dose (90 ng) of 5Z-7-Oxozeaenol diluted in 1% DMSO, or balanced salt solution containing 1% DMSO, was injected into the eyes of OIR rats at P14 at a rate of 200 nL/s using a UMP3-2 Ultra Micro Pump (World Precision Instruments, Inc. Sarasota, FL, USA). Similarly, a total of 1 μg of miR-126, miR-143, miR-145, or miR-150 mimics (GE Healthcare Dharmacon, Parramatta, NSW, Australia; Supplementary Table 2) was injected into the eye of the OIR rats at P14, and an equal amount of non-targeting scrambled negative control RNAs was injected into the contralateral eye of the same animal. At P18, the rats were sacrificed to collect the retina for further assessment. Any issues with the injection, including large backflow upon removal of the needle, hemorrhaging of external or internal vessel were noted and eyes were excluded from the study. Similarly, Brown Norway rats were intravitreally injected low dose of 5Z-7-Oxozeaenol or balanced salt solution containing 1% DMSO. The animals were then subjected to Electroretinography and Optical coherence tomography 28 days after the injection.

**Endothelial Tube Formation assay**

Endothelial tube formation was assessed as the previous described (12). BD Matrigel™ Basement Membrane Matrix (catalog no. 356234; Becton Dickinson; Bedford, MA) was thawed at 4°C overnight and then spread evenly over each well (50 μL) of 96-well plate. The plate was incubated at 37°C for 30 minutes before seeding to allow Matrigel to polymerize. HUVECs were then seeded at 2 x 10^4^ cells per well and grown in 100 μL of endothelial cell basal medium-2 (EBM-2) supplied with EGM™-2 BulletKit™ (catalog no. CC-5035; Lonza). Cells were immediately treated with 1000 nM TAK1 inhibitor (5Z-7-Oxozeaenol; catalogue no. 3604/1; Tocris Bioscience, Bristol, UK) or left untreated as controls. Images were taken for each group 6 hours after treatment. The endothelial lumen formation and branch point were quantified by using the Angiogenesis Analyzer Plugin for ImageJ version 1.48 software (http://imagej.nih.gov/ij/; provided in the public domain by the National Institutes of Health, Bethesda, MD, USA). The control samples were defined as 100% tube formation, and the percentage change in tube formation relative to control was calculated for the treated samples. Experiments were repeated at least twice.

**Cell Proliferation assay**

For the cell proliferation assay, a cell proliferation assay kit was used as per the manufacturer’s instruction (CYQUANT^®^ NF, catalog no. C35006; Life Technologies Australia). Briefly, HUVECs were seeded at 5,000 per well in 96-well plate. The cells were then treated with 5Z-7-Oxozeaenol at 1000 nM or left untreated as controls. The medium was removed and replaced with the diluted binding solution with DNA dye 24 hours or 48 hours after drug treatment. To allow the DNA being stained, the plate was incubated at 37°C for 1 hour prior to assessment of fluorescence intensity. The fluorescence intensity of samples was measured using a fluorescence microplate reader (Spark® 20M, Tecan, Switzerland) with excitation at ~485 nm and emission detection at ~530 nm. The results were normalized to the percentage of control. Experiments were repeated at least twice.

**Cell Migration**

HUVECs were seeded at a concentration of 2 x 10^5^ cells/well in a 6-well tissue culture plate prior to treatment. Cell monolayers were then wounded by scraping, followed by treatment with 5Z-7-Oxozeaenol at 1000 nM or left untreated as controls. Cells were incubated at 37 °C for 24 hours. Images of cells were taken immediately after scraping and 24 hours after 5Z-7-Oxozeaenol treatment under phase contrast microscope (TE2000-U, Nikon, Japan). For each image, the distance between the gap was quantified and normalised to the original would distance. Four fields per well were quantified with using the ImageJ version 1.48 software, and experiments were repeated at least twice.

**Aortic Ring Assay**

Aortic ring assay was performed as previously described (13). Briefly, aortae were removed from euthanized C57BL/6 mice and immediately transferred to a culture dish containing ice-cold EBM-2 supplied with EGM™-2 BulletKit™ (Lonza). The peripheral fibroadipose tissue was carefully removed. The aortae were sectioned into rings ~1 mm in width. Each aortic ring was placed in an individual well covered with 120 uL growth factor-reduced BD Matrigel™ Basement Membrane Matrix (catalog no. 356234; Becton Dickinson), and was cultured in 200uL EBM-2 for 9 days. For explants treated with 5Z-7-Oxozeaenol, 5Z-7-Oxozeaenol was added to the medium at 1000 nM. The medium was replaced and the drug was supplemented every 3 days for all groups (n=6 per group). Images of individual explants were taken on day 0 to 9 after plating using Zeiss microscope. The neo-formed vascular sprouting area was quantified with Adobe Photoshop Elements^13^.

**Electroretinography (ERG)**

A low dose (18 ng) of 5Z-7-Oxozeaenol or vehicle was injected intravitreally into the eyes of male Brown Norway rats; 28 days following the injection, the rats underwent ERG. Rats were dark-adapted overnight prior to the ERG assessment under fully dark-adapted conditions. Details for functional assessment are as previously described (14). Briefly, a pair of custom-made chloride silver active and ring-shaped reference electrodes (99.9%, A&E Metal Merchants, Sydney, NSW, AU), connected to platinum leads (F-E-30, Grass Telefactor, West Warwick, RI), was located in the central cornea and sclera, respectively. A stainless steel needle electrode (F-E2-30, Grass Telefactor) was inserted subcutaneously into the tail of the animal to replace ground. Measurements were recorded simultaneously for both eyes. ERG analysis as previously described consists of three major waveforms. Each waveform reflects the function of different retinal cells, such as the photoreceptor (a-wave), bipolar cell (b-wave), and ganglion cell (scotopic threshold response, STR).

**Spectral-domain Optical coherence tomography (OCT)**

Following ERG measurement, rat eyes were imaged using spectral domain-OCT (Envisu R2200 VHR, Bioptigen, Inc., Morrisville, NC, USA). Volume scans consisting of 1000 A-scans per 100 B-scans (Equally space across the 1.4 mm vertical dimension; 100 horizontal B-scans evenly spaced in the vertical dimension) centred on the optic nerve head (ONH; 1.4 × 1.4 × 1.57 mm) were obtained from both eyes. Four B-scans that crossed the ONH were analyzed using FIJI software (https://fiji.sc/). In each B-scan the inner limiting membrane, retinal nerve fiber layer (RNFL), inner plexiform layer and Bruch's membrane were manually segmented by a masked observer as previously described(14). Total retinal thickness (TRT) was measured from the inner limiting to Bruch’s membrane. Retinal nerve fibre layer (RNFL) thickness was measured from the inner limiting membrane to the inner aspect of the inner plexiform layer. Outer retinal thickness was measured from Bruch’s membrane to the outer plexiform layer.

**Dual-Luciferase Reporter Assay**

The oligonucleotides comprising the miR-143 target sequence were synthesized by Integrated Device Technology (Singapore), then annealed and cloned into pGL3 Luciferase Reporter Vectors (pCMV_Luciferase, Promega, Madison, WI, USA) with a reporter gene encoding firefly luciferase (pCMV_Luciferase/miRE). The miR-143 target sequence within the 3’ untranslated regions (UTRs) of TAK1 was predicted by the miRTarBase and TargetScan as described previously (15; 16). Constructs containing target sequences of miR-143 (Supplementary Table 3) was confirmed by restriction enzyme digestion and Sanger sequencing at AGRF (Melbourne, VIC, Australia). Transfection was performed in 12-well plates with 80% confluent HEK293A cells. Two hundred nanogram of firefly luciferase plasmid and 50 ng of the Renilla luciferase plasmid (pRL-SV40; Promega) were co-transfected into cells by using Lipofectamine 2000 (catalog no. 11668027; Life Technologies Australia). Cells were simultaneously co-transfected with 40 nM miR-143 mimics or scrambled negative control RNAs. For each transfection assay, the pCMV_Luciferase vector serves as positive control. After 48 hours, cell lysates were prepared from each transfected culture, and Dual-Luciferase assay was performed using the dual luciferase kit (catalog no. E1910; Promega). Luminescence was measured by using a microplate reader (Spark® 20M, Tecan). Luciferase activity was normalized to Renilla luciferase activity and to positive controls.

**References**

1. Nixon B, Stanger SJ, Mihalas BP, Reilly JN, Anderson AL, Dun MD, Tyagi S, Holt JE, McLaughlin EA: Next Generation Sequencing Analysis Reveals Segmental Patterns of microRNA Expression in Mouse Epididymal Epithelial Cells. PloS one 2015;10:e0135605

2. Anderson AL, Stanger SJ, Mihalas BP, Tyagi S, Holt JE, McLaughlin EA, Nixon B: Assessment of microRNA expression in mouse epididymal epithelial cells and spermatozoa by next generation sequencing. Genomics data 2015;6:208-211

3. Friedlander MR, Chen W, Adamidi C, Maaskola J, Einspanier R, Knespel S, Rajewsky N: Discovering microRNAs from deep sequencing data using miRDeep. Nature biotechnology 2008;26:407-415

4. Andrews S: FastQC: a quality control tool for high throughput sequence data. 2010;

5. Trapnell C, Pachter L, Salzberg SL: TopHat: discovering splice junctions with RNA-Seq. Bioinformatics 2009;25:1105-1111

6. Trapnell C, Williams BA, Pertea G, Mortazavi A, Kwan G, van Baren MJ, Salzberg SL, Wold BJ, Pachter L: Transcript assembly and quantification by RNA-Seq reveals unannotated transcripts and isoform switching during cell differentiation. Nat Biotechnol 2010;28:511-515

7. Trapnell C, Roberts A, Goff L, Pertea G, Kim D, Kelley DR, Pimentel H, Salzberg SL, Rinn JL, Pachter L: Differential gene and transcript expression analysis of RNA-seq experiments with TopHat and Cufflinks. Nat Protoc 2012;7:562-578

8. Metsalu T, Vilo J: ClustVis: a web tool for visualizing clustering of multivariate data using Principal Component Analysis and heatmap. Nucleic Acids Res 2015;43:W566-570

9. Livak KJ, Schmittgen TD: Analysis of relative gene expression data using real-time quantitative PCR and the 2(-Delta Delta C(T)) Method. Methods 2001;25:402-408

10. Hewing NJ, Weskamp G, Vermaat J, Farage E, Glomski K, Swendeman S, Chan RV, Chiang MF, Khokha R, Anand-Apte B, Blobel CP: Intravitreal injection of TIMP3 or the EGFR inhibitor erlotinib offers protection from oxygen-induced retinopathy in mice. Invest Ophthalmol Vis Sci 2013;54:864-870

11. Chen J, Connor KM, Aderman CM, Willett KL, Aspegren OP, Smith LE: Suppression of retinal neovascularization by erythropoietin siRNA in a mouse model of proliferative retinopathy. Invest Ophthalmol Vis Sci 2009;50:1329-1335

12. Donovan D, Brown NJ, Bishop ET, Lewis CE: Comparison of three in vitro human 'angiogenesis' assays with capillaries formed in vivo. Angiogenesis 2001;4:113-121

13. Baker M, Robinson SD, Lechertier T, Barber PR, Tavora B, D'Amico G, Jones DT, Vojnovic B, Hodivala-Dilke K: Use of the mouse aortic ring assay to study angiogenesis. Nat Protoc 2012;7:89-104

14. Zhao D, Nguyen CT, Wong VH, Lim JK, He Z, Jobling AI, Fletcher EL, Chinnery HR, Vingrys AJ, Bui BV: Characterization of the Circumlimbal Suture Model of Chronic IOP Elevation in Mice and Assessment of Changes in Gene Expression of Stretch Sensitive Channels. Front Neurosci 2017;11:41

15. Hsu JB, Chiu CM, Hsu SD, Huang WY, Chien CH, Lee TY, Huang HD: miRTar: an integrated system for identifying miRNA-target interactions in human. BMC bioinformatics 2011;12:300

16. Chen X, Rosbash M: mir-276a strengthens Drosophila circadian rhythms by regulating timeless expression. Proceedings of the National Academy of Sciences of the United States of America 2016;113:E2965-2972

**Supplemental Material- Supplementary Figures**

**Supplementary Figure 1**

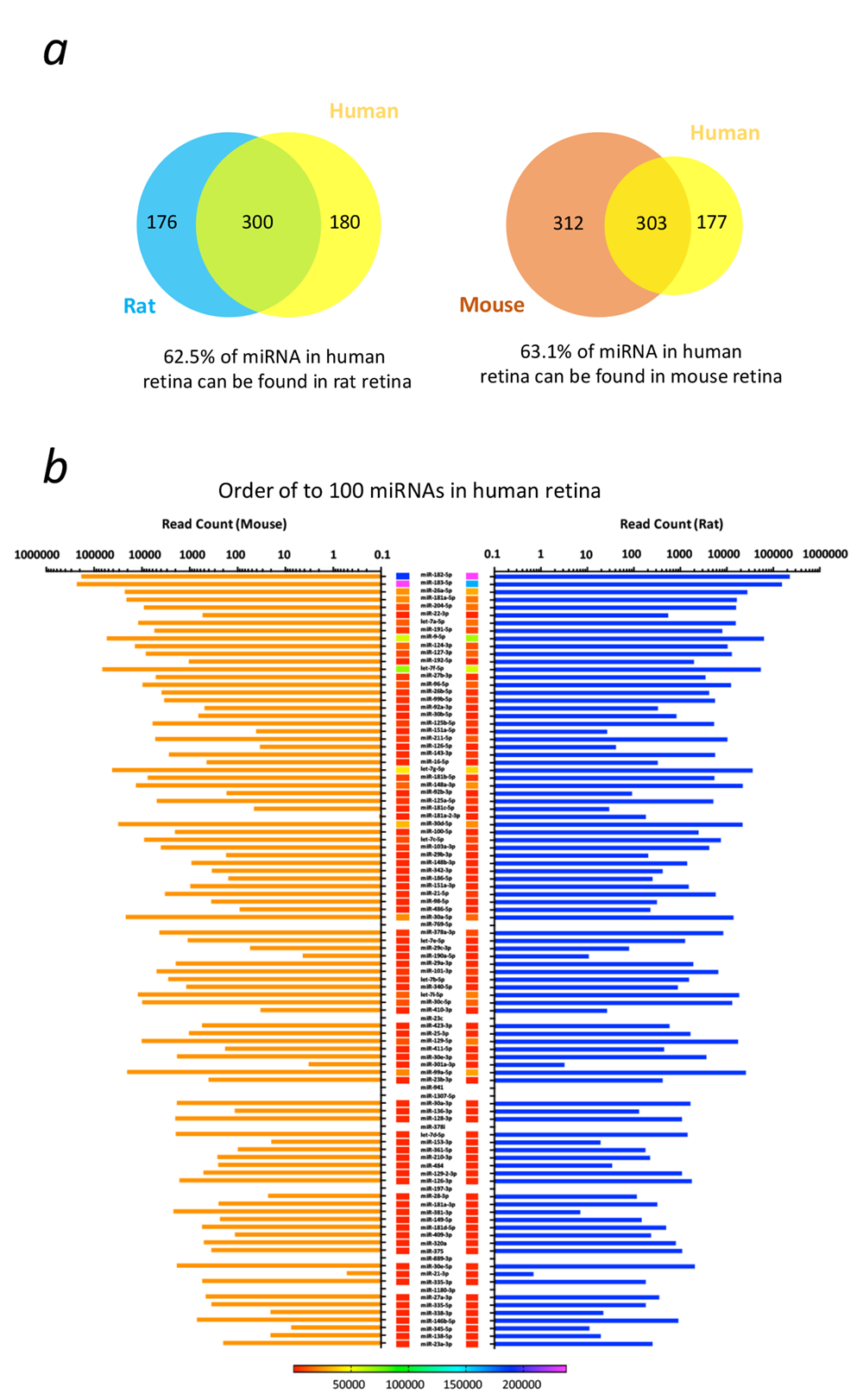

**Supplementary Figure 1. Rat and mouse have highly homologous miRNA expression in the retina.** **a)** The venn diagrams showed that rat and mouse both share similar retinal miRNA expression profile with humans. **b)** The retinal miRNA from both rat and mouse were extracted and subjected to miRNA-Seq. The expression of retinal miRNAs in either rat or mouse were aligned and matched to the top 100 miRNAs expressed in the human retina. The mean read count of each miRNAs from miRNA-Seq was depicted in the diagram (n=3 per group).

**Supplementary Figure 2**

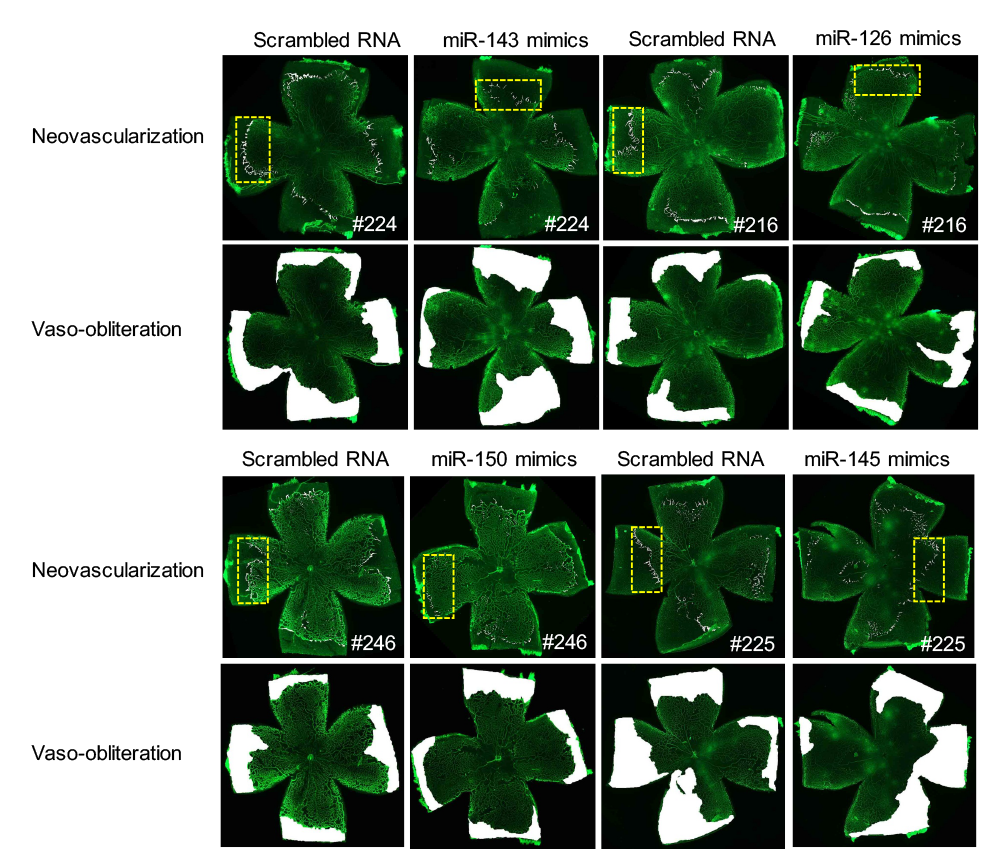

**Supplementary Figure 2. The uncropped images of the retinal flat-mount used in Figure 3b.** The area with the dashed line was snipped as a representative for retinal neovascularization. The whitish area represents the avascular area in the retina.

**Supplementary Figure 3**

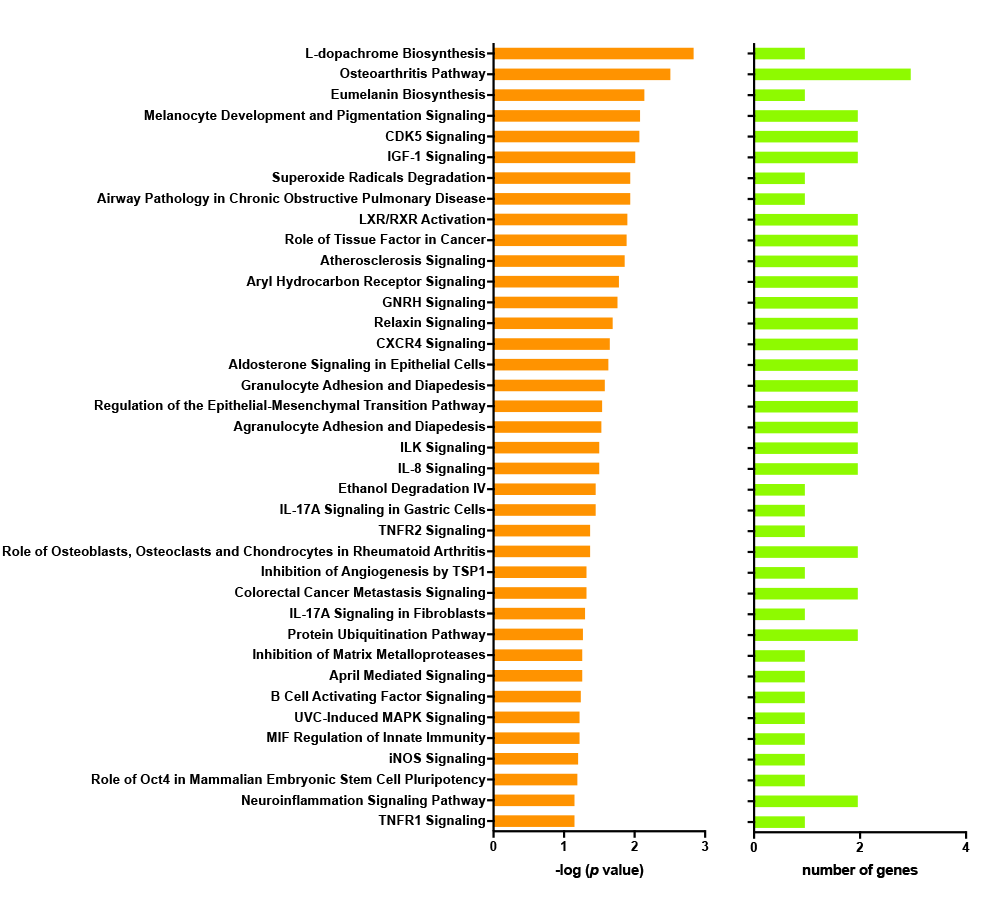

**Supplementary Figure 3.** A range of canonical pathways was involved in the suppression of retinal neovascularization in the OIR rats by miR-143 mimics treatment.

**Supplementary Figure 4**

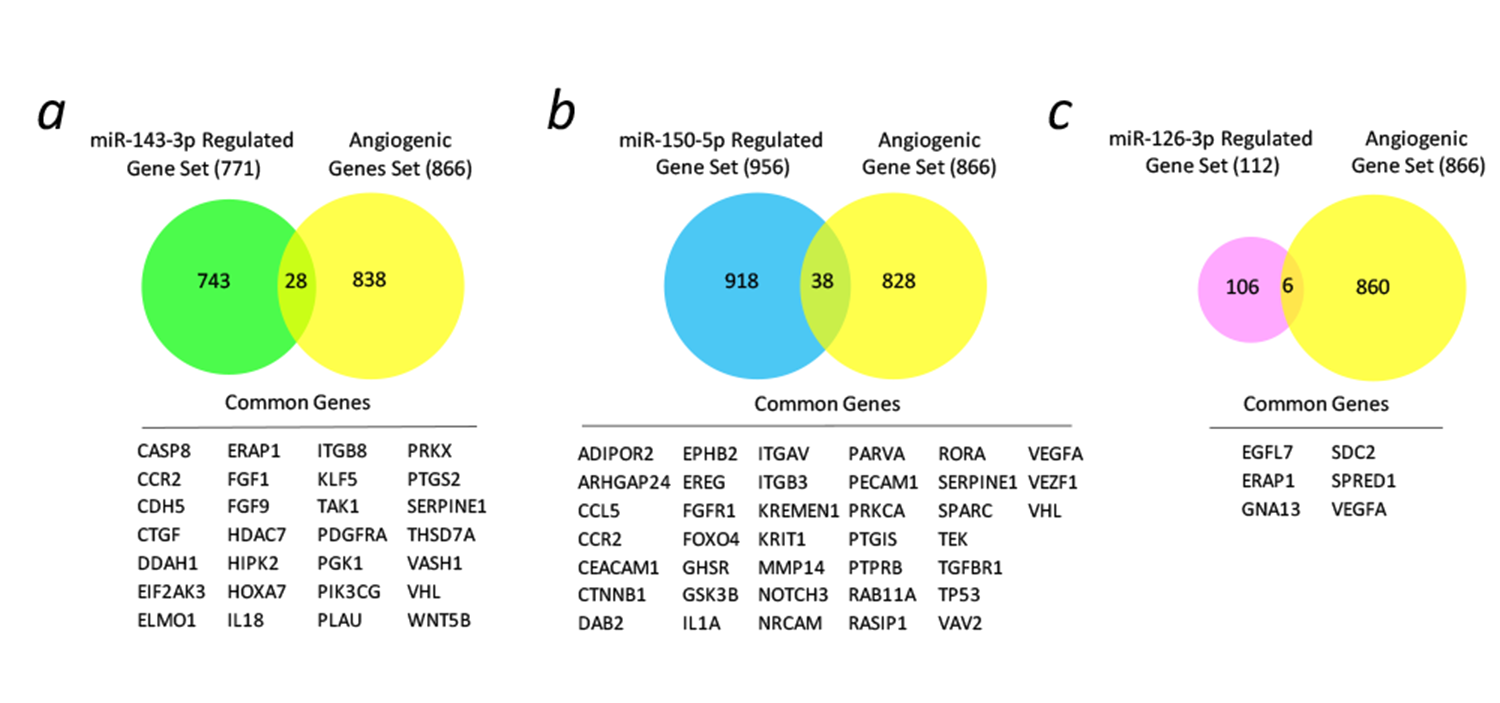

**Supplementary Figure 4. Putative pro-angiogenic genes that regulated by miR-143 or miR-150 or miR-126 were identified.** **a)** The venn diagram depicted the overlap between the genes regulated by miR-143 and pro-angiogenic gene sets. 28 genes were identified. **b)** The venn diagram depicted the overlap between the genes regulated by miR-150 and pro-angiogenic gene sets. 38 genes were identified. **c)** The venn diagram depicted the overlap between the genes regulated by miR-150 and pro-angiogenic gene sets. 6 genes were identified.

**Supplementary Figure 5**

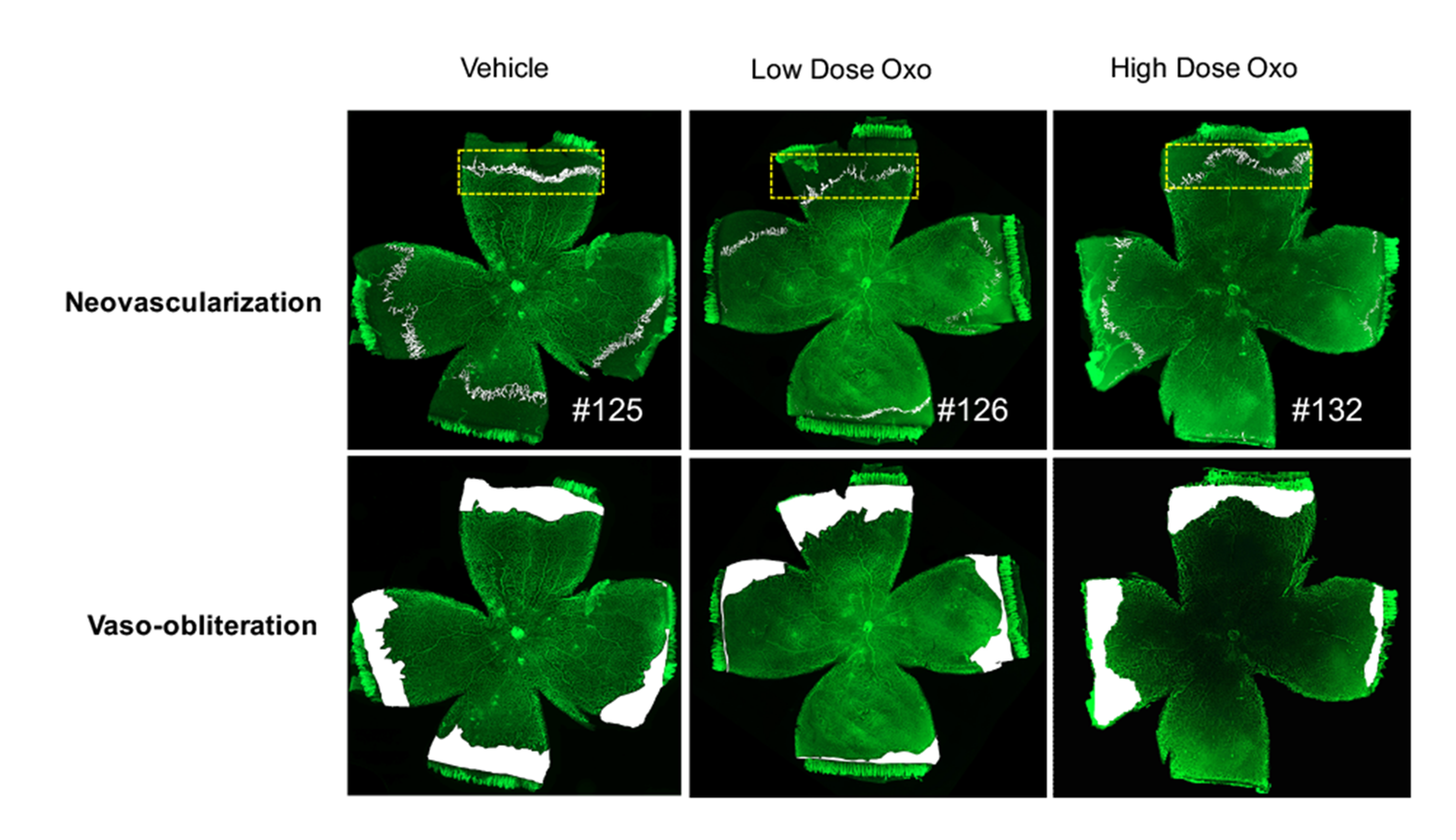

**Supplementary Figure 5. The uncropped images of the retinal flat-mount used in Figure 4g.** The area with the dashed line was snipped as a representative for retinal neovascularization. The whitish area represents the avascular area in the retina. Oxo: 5Z-7-Oxozeaenol.

**Supplemental Material- Supplementary Tables**

**Supplementary Table 1. TaqMan probe sequences used for real-time quantitative polymerase chain reaction**

| **Assay Name** | **Source** | **Assay ID** | **TaqMan probe sequence** |
| --- | --- | --- | --- |
| rno-miR-143 | Applied Biosystems | Rn03466026_pri | GCGGAGCGCCUGUCUCCCAGCCUGAGGUGCAGUGCUGCAUCUCUGGUCAGUUGGGAGUCUGAGAUGAAGCACUGUAGCUCAGGAAGGGAGAAGAUGUUCUGCAGC |
| rno-miR-150 | Applied Biosystems | Rn03466044_pri | CUUCUCAAGGCCCUGUCUCCCAACCCUUGUACCAGUGCUGUGCCUCAGACCCUGGUACAGGCCUGGGGGACAGGGACUUGGGGAC |
| rno-miR-126 | Applied Biosystems | Rn03464585_pri | UGACAGCACAUUAUUACUUUUGGUACGCGCUGUGACACUUCAAACUCGUACCGUGAGUAAUAAUGCGUGGUCA |
| rno-miR-145 | Applied Biosystems | Rn03466035_pri | CACCUUGUCCUCACGGUCCAGUUUUCCCAGGAAUCCCUUGGAUGCUAAGAUGGGGAUUCCUGGAAAUACUGUUCUUGAGGUCAUGGCU |
| rno-miR-451 | Applied Biosystems | Rn03465141_pri | UUUGGGAAUGGCGAGGAAACCGUUACCAUUACUGAGUUUAGUAAUGGUAAUGGUUCUCUUGCUGCUCCCACA |
| U6 snRNA | Applied Biosystems | 001973 | GTGCTCGCTTCGGCAGCACATATACTAAAATTGGAACGATACAGAGAAGATTAGCATGGCCCCTGCGCAAGGATGACACGCAAATTCGTGAAGCGTTCCATATTTT |
| Rat Actb | Applied Biosystems | Rn00667869_m1 | N/A |
| Rat TAK1 | Applied Biosystems | Rn01437015_m1 | N/A |
| Rat TNF-α | Applied Biosystems | Rn01525859_g1 | N/A |
| Rat IL-6 | Applied Biosystems | Rn01410330_m1 | N/A |
| Rat TGF-β1 | Applied Biosystems | Rn00572010 | N/A |
| Rat VEGF-A | Applied Biosystems | Rn01511602_m1 | N/A |
| Rat VEGF-B | Applied Biosystems | Rn01454585_g1 | N/A |
| Rat VEGF-C | Applied Biosystems | Rn01488076_m1 | N/A |

**Supplementary Table 2. Oligonucleotide sequences of miRNA mimic**

| **miRNA mimics** | **Source** | **Catalogue number** | **Sequences (5’-3’)** |
| --- | --- | --- | --- |
| rno-miR-143-3p | Dharmacon | MIMAT0000849 | UGAGAUGAAGCACUGUAGCUCA |
| rno-miR-150-5p | Dharmacon | MIMAT0000853 | UCUCCCAACCCUUGUACCAGUG |
| rno-miR-126a-3p | Dharmacon | MIMAT0000832 | UCGUACCGUGAGUAAUAAUGCG |
| rno-miR-145-5p | Dharmacon | MIMAT0000851 | GUCCAGUUUUCCCAGGAAUCCCU |
| Scrambled RNA | Dharmacon | MIMAT0000039 | UCACAACCUCCUAGAAAGAGUAGA |

**Supplementary Table 3. The expression level of down-regulated retinal miRNAs in the OIR rats at P14 was restored at P29.**

| **MicroRNA** | **OIR (P14)** | **OIR (P29)** | **Log_2_ (Fold Change)** | **Adjusted p value** | **Expression Level** |
| --- | --- | --- | --- | --- | --- |
| rno-miR-150-5p | 84.00 ± 20.66 | 520.33 ± 18.56 | 2.47240 | 9.54E-06 | up |
| rno-miR-143-3p | 5443.67 ± 1962.62 | 21591.00 ± 1240.32 | 1.86802 | 3.85E-05 | up |
| rno-miR-451-5p | 15.67 ± 2.89 | 152.00 ± 30.51 | 3.05133 | 8.40E-05 | up |
| rno-miR-126a-3p | 595.67 ± 190.42 | 1647.00 ± 257.49 | 1.32408 | 2.88E-04 | up |
| rno-miR-145-5p | 66.00 ± 34.22 | 201.00 ± 32.51 | 1.52989 | 1.98E-03 | up |

**Supplementary Table 4. Oligonucleotide sequences for the luciferase reporter constructs.**

| **Gene** | **Seed position** | **Sequences (5’-3’)** |
| --- | --- | --- |
| TAK1 | 386-406 | GGCCGCATGACTTTATTCTTGTATCTCATCTCAAAATATTAATAATTTTTTG |
